# In Troyer syndrome Spartin loss induces Complex I impairments and alters pyruvate metabolism

**DOI:** 10.1101/488239

**Authors:** Chiara Diquigiovanni, Christian Bergamini, Rebeca Diaz, Irene Liparulo, Francesca Bianco, Luca Masin, Antonia Tranchina, Francesco Buscherini, Titia Anita Wischmeijer, Tommaso Pippucci, Emanuela Scarano, Duccio Maria Cordelli, Romana Fato, John Milton Lucocq, Marco Seri, Silvia Paracchini, Elena Bonora

## Abstract

Growth delay and retardation are complex phenotypes which can results by a range of factors including genetics variants. We identified a novel homozygous frameshift mutation, c.892dupA, in *SPART* gene, in two brothers with short stature and psychomotor retardation, born from healthy consanguineous parents. Mutations in *SPART* are the cause of Troyer syndrome, an autosomal recessive form of spastic paraplegia resulting in muscle weakness, short stature and cognitive defects. *SPART* encodes for Spartin, a protein with different cellular functions, such as endosomal trafficking and mitochondrial stability.

We evaluated the effects of Spartin loss by transiently silencing *SPART* in human neural stem cells (hNSCs) and by generating an SH-SY5Y cell line model carrying the c.892dupA mutation via CRISPR/Cas9. In both models, we observed an altered neuronal growth and an increase in neurite outgrowth. In the SH-SY5Y cell line carrying the c.892dupA mutation, Spartin absence led to an altered distribution of mitochondria, and to a severe decrease in the NADH-dehydrogenase activity of mitochondrial Complex I. These impairments determined an energetic failure with a decrease in ATP synthesis due to a halt in mitochondrial oxidative phosphorylation, increased reactive oxygen species production, and alteration in intracellular Ca^2+^ homeostasis. Transient re-expression of Spartin in mutant cells restored an intracellular Ca^2+^ level. Mutant cells presented a significant increase in extracellular pyruvate, which may result from increased glycolysis due to impaired Complex I activity. Consistently, Spartin loss led to an over-activation of Signal Transducer and Activator of Transcription 3 (STAT3) factor, a key regulator of glycolysis.

These data demonstrate that Spartin loss leads to a profound bioenergetics imbalance with defective OXPHOS activity, and this altered metabolism might underlie Troyer syndrome and neurodevelopmental delays.

## Introduction

Neurodevelopmental disorders affect 2–5% of individuals and are genetically heterogeneous (DDS Deciphering Consortium, 2017). They constitute a large proportion of the life-long global health burden in terms of medical care, hospitalizations, and mortality (Khokha *et al.*, 2017). Next generation sequencing (NGS) technologies have proved to be a powerful tool for the identification of rare mutations that cause these disorders (Short *et al.*, 2018; Wright *et al.*, 2018; Vissers and Veltman, 2015; Firth and Wright, 2011). For example, recurrent epimutations at chromosome 11 or unidisomy of chromosome 7 have been implicated in growth retardation. But in most cases the genetic cause remains unknown. The identification of a causative mutation can support diagnosis, prognosis, and available treatment (Seaver and Irons, 2009). We evaluated two brothers born from healthy consanguineous parents (first degree cousins), referred for pre- and post-natal growth retardation, syndromic short stature and developmental delay with severe speech impairment, for whom no genetic diagnosis had been identified. We performed a whole exome sequencing analysis in the two brothers and identified a novel homozygous mutation in the *SPART* gene, c.892dupA, inserting a premature stop codon in the Spartin protein, where loss-of-function mutations have been found in Troyer syndrome (OMIM #275900). Although no suspect of Troyer syndrome had been hypothesized for the two affected sibs, a careful re-evaluation identified overlapping features with this disorder, a rare autosomal-recessive form of spastic paraplegia, characterized by lower extremity spasticity and weakness, short stature, cognitive defects and distal amyotrophy caused by the degeneration of corticospinal tract axons (Alazami *et al.*, 2015; Tawamie *et al.*, 2015; Manzini *et al.*, 2010; Patel *et al.*, 2002). *S*partin is a multifunctional protein consisting of a N-terminal Microtubule Interacting and Trafficking (MIT) domain and a C-terminal senescence domain (Bakowska *et al.*, 2005; Ciccarelli *et al.*, 2003) (Supplementary Fig. 1A). Spartin is involved in different cellular processes such as the intracellular trafficking of the epidermal growth factor receptor and the turnover of lipid droplets, bone morphogenetic protein (BMP) signalling and cytokinesis (Renvoise *et al*., 2010; Eastman *et al.*, 2009; Tsang et al., 2009; Edwards *et al.*, 2009; Bakowska *et al*., 2007). In particular, *Spg20*-/- mice showed a prominent number of binucleated chondrocytes in epiphyseal growth plates of bones, indicating that impaired cytokinesis during bone formation is related to the short stature and skeletal defects observed in Troyer syndrome (Renvoisé *et al.*, 2012).

Spartin expression has been identified in different tissues at embryonic and adult stages. In the Eurexpress mouse database (http://www.eurexpress.org) expression of the homologous murine *Spg20* was identified in the nervous and olfactory systems of the developing mouse at embryonic day 14.5 (Diez-Roux *et al.*, 2011). In the Human Protein Atlas *SPART* expression is specific for neuronal cells in adult cortex (https://www.proteinatlas.org/ENSG00000133104-SPG20).

A few studies suggested that Spartin loss might impair mitochondrial stability, since absence of Spartin induced an alteration in the mitochondrial network and a decrease in the mitochondrial membrane potential (Milewska and Byrne, 2015; Truong *et al.*, 2015; Joshi and Bakowska, 2011), but a thorough analysis of mitochondrial respiration and OXPHOS activity has not been carried out yet. Our aim was to gain further insight into the role of Spartin on mitochondrial activities, investigating the effects of the novel *SPART* loss-of-function mutation. We first evaluated the effects of Spartin loss by gene silencing in human neural stem cells (hNSCs). Then, we characterized the *SPART* mutation found in the two affected sibs by generating a CRISPR/Cas9- modified neuroblastoma-derived SH-SY5Y cell line. Mutant cells developed an altered neuronal growth, exhibited significantly longer neurites, associated to an altered distribution and structure of the mitochondrial network. In terms of cell metabolism, we found a severe decrease in Complex I activity coupled to an increased production of mitochondrial reactive oxygen species (ROS) and identified an elevated extracellular pyruvate, indicative of a defective mitochondrial oxidation of this molecule. Our data suggest that the mitochondrial impairment could affect neuronal by inducing an energetic failure coupled to excessive ROS production and promoting an extended axonal morphology prone to neurodegeneration. These data explain the pleiotropic effects of *SPART* mutations on biological pathways underlying Troyer syndrome and that might be relevant for other form of neurodevelopmental conditions.

## Patients and method

### Subjects

Two brothers born from consanguineous healthy parents, first-degree cousins of Moroccan origin, were first evaluated at the Clinical Genetics Unit when they were 42-months and 12-months-old, respectively. Family history was unremarkable. They both presented with IUGR (intrauterine growth restriction), stature and weight below -2 SD (standard deviations), relative *macrocrania*, dysmorphic features (very long eyelashes, dolichocephaly, prominent maxilla, *pectus excavatum*) and mild psychomotor retardation, with severe language delay. They started walking independently at 18 months. Other shared anomalies included delayed bone age, *pes planus*, euphoric behavior, and joint hyperlaxity. The first evaluation did not disclose signs of neuromuscular involvement. Genetic analyses included: karyotype and analysis of subtelomeric regions, UPD7 and H19 methylation analysis, and mutation screening of *PNPLA6* (OMIM#603197), all of which were negative.

After a 5 years follow-up, the eldest brother (age 8 years and 9 months) gradually developed muscular hypotrophy in upper and lower limbs, increased muscle tone in the lower limbs (distal>proximal) and brisk deep tendon reflexes. Sporadic aggressive behavior and inappropriate crying was reported by parents. The younger sib (6 years and 3 months) developed muscular hypotrophy as well, mild hyper-reflexia and difficulty to walk on toes or heels. No cerebellar signs were reported, nor dysarthria/tongue dyspraxia but language impairment remained severe. Stature was constantly around the 3^rd^ percentile in both sibs. The eldest brother showed a partial Growth Hormone (GH) deficiency, treated with a GH analogue.

Detailed Materials and Methods are available in Supplementary material section.

## Results

### Whole exome sequencing identified a novel loss-of-function mutation in *SPART* gene

We performed a combined analysis of high-density SNPs genotyping and WES in the two affected brothers (II-1; II-2, Fig. 1A) born in a consanguineous family. The two siblings presented with history of intrauterine growth restriction, stature and weight below -2 SD, relative macrocrania, dysmorphic features (very long eyelashes, dolichocephaly, prominent maxilla, *pectus excavatum*, Fig. 1B) and psychomotor retardation, with severe language delay (for an exhaustive description of cases see the Materials and Methods section). Considering the degree of inbreeding, the search for runs of homozygosity (ROH) using SNP data from Illumina 350K array identified a region of homozygosity on chromosome 13 (5 Mb). WES analysis identified a novel insertion on chr13:g3690561insT (hg19), leading to a c.892dupA (NM_001142294) in the *SPART* gene (OMIM *607111). The mutation was homozygous in the two affected sibs and caused a frameshift with the insertion of a premature stop codon (p.Thr298Asnfs*17) in the protein Spartin. The variant was carried by the parents (Figure 1A, I-1 and I-2) and was not present in the Exome Aggregation database (ExAc) and Genome Aggregation database (gnomAD) (as accessed on 20/11/2018), nor in in-house internal whole exome database of 650 exomes.

**Figure 1.**
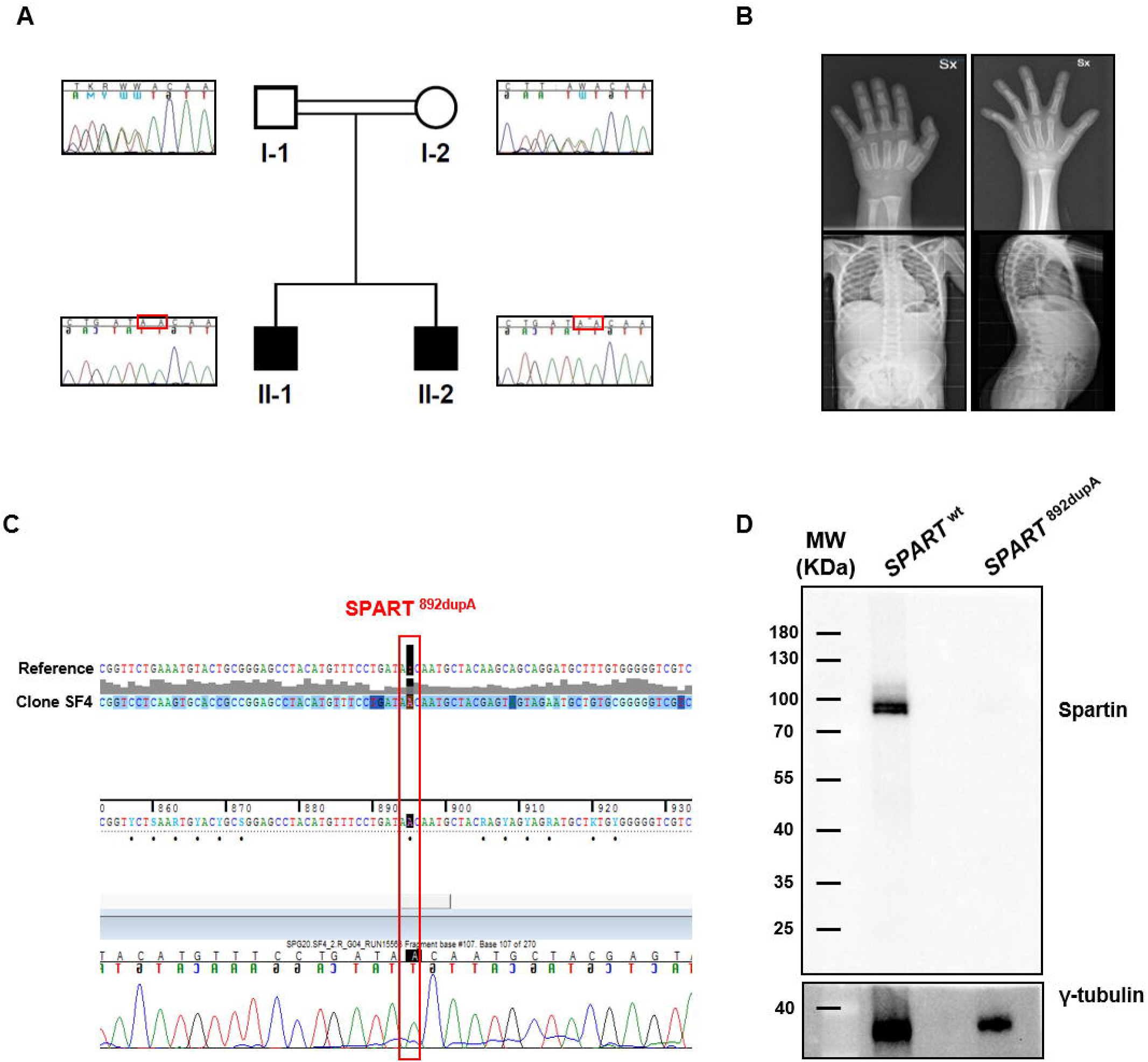
Identification of *SPART* c.892dupA variant. **(A)** Pedigree of the consanguineous family and electropherograms of the sequences in family members showing the co-segregation of the change with the spastic paraplegia phenotype. The two sibs are homozygous for the mutation, whereas both parents are heterozygous carriers. **(B)** Representative images of patient’s skeletal defects. Hand X-ray showed a delayed bone age, 1 year at 2 years of chronological age and 2.5 years at 5.5 years of chronological age. **(C)** Generation of *SPART* c.892dupA knock-in SH-SY5Y cell line. Electropherogram of the SH-SY5Y clone sequence carrying the mutation c.892dupA in *SPART* gene and alignment between reference sequence and clone sequence are reported. **(D)** Representative western blot of Spartin protein from *SPART*^wt^ and *SPART*^892dupA^-SH-SY5Y clone using a specific anti-Spartin antibody (13791-1-AP, N-terminal, catalog number AG815; ProteinTech). Western confirmed that in *SPART*^892dupA^ cells, the predicted 33KDa mutant Spartin is not produced, whereas in control cells Spartin is normally synthesized and migrated at the expected molecular weight of 75-84 KDa. Gamma tubulin was used as endogenous control.

### Generation of biological models

To understand the effect of *SPART* ablation we generated two different biological models. First, we transiently silenced the gene in human Neural Stem cells (hNSCs), using siRNA specific to *SPART* transcripts *vs* scramble-treated cells. Silencing efficiency was evaluated via western blot analysis (Supplementary Fig. 1B). Then, we generated a stable SH-SY5Y cell line, knocking-in the c.892dupA mutation using the CRISPR/Cas9 technology, in order to study the effects of this variant in a stable neuronal model. Cells were transfected with gRNA and Cas9-nickase plasmids and the oligo DNA carrying the c.892dupA variant to insert the specific modification in the SH-SY5Y genome (Fig. 1C). Protein expression was assessed on the generated clones. The variant is predicted to generate a shorter protein of 33KDa, however western blot analysis showed that Spartin in *SPART*^892dupA^ cells was completely absent (Fig. 1D), probably because of nonsense-mediated mRNA decay mechanisms (Brogna and Wen, 2009).

### Spartin depletion affects neuronal morphology leading to neuronal differentiation

To understand the effect of *SPART* loss on neuronal morphology, we transiently silenced the gene in human Neural Stem cells (hNSCs) up to 8 days, using siRNA specific to *SPART* transcripts *vs* scramble-treated cells (Fig. 2A, B). In *SPART*-silenced hNSCs, we observed an increased neurite outgrowth, visualized via nestin staining (Fig. 2A, panel f and Fig. 2B, panel a) compared to scramble-treated hNSCs (Fig. 2A, panel e and Fig. 2B, panel b). According to this finding, in the stable knock-in cell line carrying the c.892dupA mutation, *SPART*^892dupA^, immunostaining for PGP9.5, a specific neuronal marker localized to cell bodies and neurites, revealed an altered neuronal morphology compared to control *SPART*^wt^ cells (Fig. 2C). *SPART^8^*^92dupA^ cells showed extensive and branched neurite-like formations (Fig. 2C, panels g, h) compared to *SPART*^wt^ (Fig. 2C, panels c, d). Neuritogenesis was quantified by performing a quantitative evaluation of the number and length of neurites with the NeuronGrowth software. *SPART^8^*^92dupA^ cells showed significantly longer neuronal processes, compared to *SPART*^wt^ cells (p=0.0001; Supplementary Fig. 1C) and an increase in average number of neurites per cell extending from the cell body (p=0.0337; Supplementary Fig. 1D).

**Figure 2.**
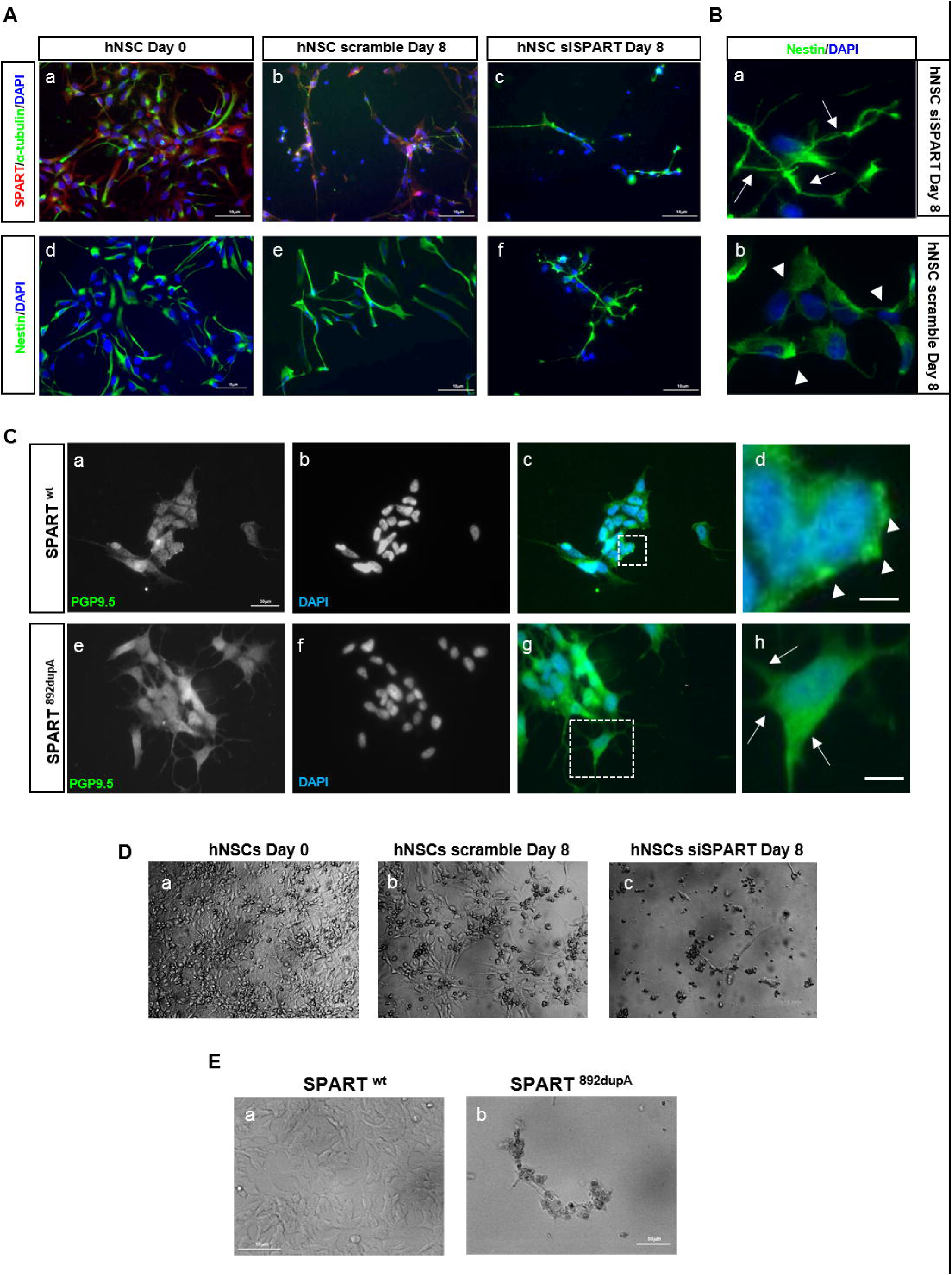
Spartin depletion affects neuronal morphology and cells growth. **(A)** Representative immunofluorescence images showing *SPART*-silencing in hNSCs. Images showed hNSCs day 0 (not silenced, panels a, d), hNSC scramble day 8 (panels b, e) and hNSC *SPART*- silenced (panels c, f). Panel a-c showed immunostaining for Spartin (red), α-tubulin (green) and DAPI (blue). Panels d-f showed immunostaining for Nestin (green) and DAPI (blue). Images showed an increased neuronal outgrowth in Spartin-depleted hNSCs compared to controls. Scale bars 10 µm. **(B)** Photograms in panel a and b are magnifications showing respectively hNSC *SPART*-silenced and hNSC scramble cells at day 8, immunostained for Nestin (green) and DAPI (blue). Arrows indicate neurite extensions that are absent in scramble-treated cells (arrowheads). Scale bars 50 µm. **(C)** Representative immunofluorescence images of *SPART*^wt^ and *SPART*^892dupA^SH-SY5H cell lines stained against the neuronal marker PGP9.5. *SPART*^892dupA^ cells showed an increased neuronal morphology compared to *SPART*^wt^ cells. Scale bars 50µm. Merged images (panels d and h) show at higher magnification the insets of the regions in panels c and g, respectively. Arrowheads in panel d showed the absence of neurites formation in *SPART*^wt^ cells. Arrows in panel h indicate the neurite length generation in *SPART*^892dupA^ cell line. Scale bar 50 µm. **(D)** Representative images of hNSCs after 8 days of *SPART* depletion (panel c, hNSCs si*SPART* Day 8), showing a strong cell loss compared to control cells (panel a and b, hNSCs at day 0 and hNSC scramble-transfected at day 8). Scale bars 10 µm. **(E)** Representative images showing *SPART*^wt^ (control) and *SPART*^892dupA^ cell lines, showing a strong cell loss in *SPART*^892dupA^ cells (panel b) compared to control cells (panel a). Scale bars 50µm.

Moreover, observation of the cell culture revealed that both cell models were characterized by an extensive cell loss in Spartin-depleted cells compared to controls (Fig. 2D-E). These data indicate that the novel variant leads to an increased neurite outgrowth and defective cell growth/survival.

### Spartin loss alters cell morphology in 3D cultures

Since *SPART*^892dupA^ cells showed an increased neuronal differentiation, we evaluated cell growth and morphology in a matrix-free environment. When cells were shifted from a 2D to a 3D micro environment (spheroids), *SPART*^wt^ and *SPART*^892dupA^ cultures showed a different morphology. Twenty-four hours after seeding both cell lines started to aggregate, and they grew into spheroids like structure at 72 hours (Fig. 3A). *SPART*^wt^ cells formed condensed and disorganized aggregates (Fig. 3A, panels b, c), whereas *SPART*^892dupA^ cells formed rounder and more compact aggregated more similar to spheroids (Fig. 3A, panels e, f). These differences started to be detectable at 24 hours but became clear at 72 hours. According to the classification by Kenny et al. (Kenny *et al.*, 2007), *SPART*^wt^ derived spheroids could be classified as a “grape-like” group, characterized by cancer stem cell-like properties, whereas *SPART*^892dupA^-derived spheroids could be classified as “round group”, characteristic of non-malignant and more differentiated cells (Fig. 3B).

**Figure 3.**
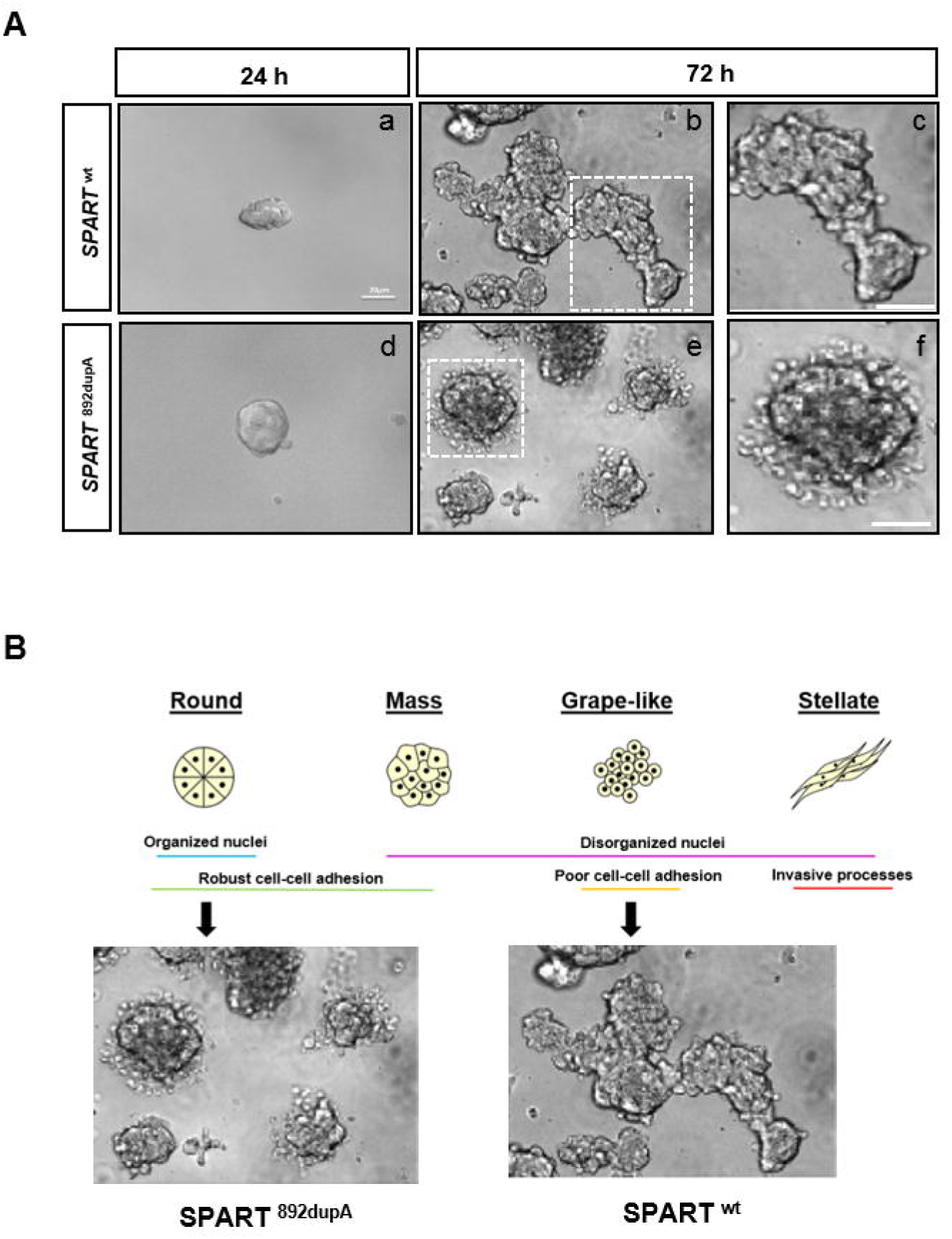
Morphological characterization of *SPART*^892dupA^ spheroids. **(A)** Morphological evaluation of *SPART*^wt^ and *SPART*^892dupA^-derived spheroids at 24 and 72 hours (panels a, b representing *SPART*^wt^ spheres and panels d, e representing *SPART* ^892dupA^ spheres). Scale bar 20 µm. Both wild-type and mutant cell lines aggregate at 24 hours and formed spheroids after 72 hours. Panels c and f showed magnification of the indicated regions in panels b and e. *SPART*^892dupA^ spheroids were more rounded (panel f) than control *SPART*^wt^ cells, whereas (panel e) formed more condensed and disorganized aggregates. **(B)** Characterization of distinct morphological groups in 3D culture. Cells could be classified into four groups (Round, Mass, Grape-like and Stellate). Schematic description for each morphological group is reported according to Kenny et al., 2007. *SPART*^wt^-derived spheroids could be classified as a “grape-like” group, whereas *SPART*^892dupA^- derived spheroids could be classified as “round group”, characteristic of more differentiated cells.

### Spartin loss alters the mitochondrial network

Previous studies showed that Spartin co-localized with mitochondria and contributed to mitochondrial stability (Lu *et al*, 2006; Milewska and Byrne, 2015; Joshi and Bakowska, 2011). Therefore, we evaluated the effects of the c.892dupA mutation on different mitochondrial characteristics in mutant *SPART*^892dupA^ and control *SPART*^wt^. Assessment of mitochondrial network was performed by live-cell microscopy in mitoGFP-transfected cells (Fig. 4A). The total mitochondrial mass was similar between *SPART*^wt^ and *SPART*^892dupA^ cells (Supplementary Fig. 2A), but mitochondria in *SPART*^892dupA^ cells exhibited a decreased interconnectivity, indicated by an elevated area-to-perimeter ratio, compared to the *SPART*^wt^ control cells (p<0.0001; Fig. 4B). Furthermore, a decrease in the number of mitochondria was observed in neurite-like extensions in mutants compared to controls (Fig. 4A, arrows). These data supported the hypothesis that Spartin loss could cause mitochondrial mobility impairments.

**Figure 4.**
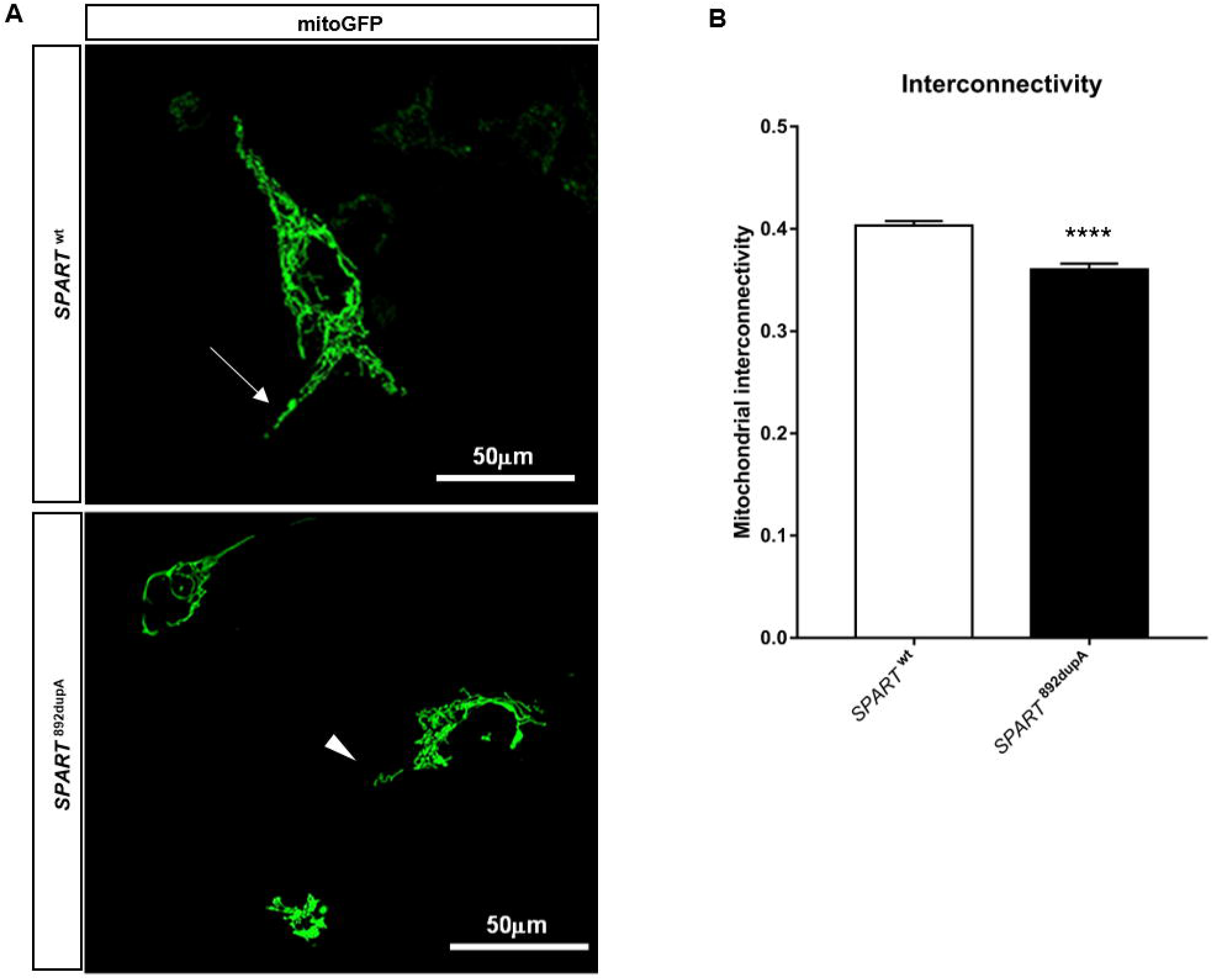
Spartin loss alters the mitochondrial morphology. **(A)** Representative z-stack image of mitochondrial network evaluated using mito-GFP probe by live cell imaging. Control cells showed a more diffused distribution of mitochondria within the cytoplasm of cells, also in neurites (upper panel, arrows). However, *SPART*^892dupA^ cells showed a notable perinuclear distribution and absence of mitochondria in neurites (lower panel, arrowheads). Scale bars 50µm. **(B)** Quantitative assessment of mitochondrial interconnectivity measured via live cell imaging (n=35 live cells measured for each cell line) using *ImageJ* Mitochondrial Morphology plugin. We observed the presence of mitochondria in neurite-like extension in *SPART*^wt^ cells (arrow) that are absent in *SPART*^892dupA^ (arrowhead) \**P* < 0.05, ****P < 0.0001, means ± SEM.

### Spartin loss determines a specific OXPHOS Complex I impairment

Mitochondrial morphology is related to its functionality (Mishra and Chan 2016), therefore we measured the oxygen consumption rates in *SPART*^wt^ and *SPART*^892dupA^ intact cells, in absence and in presence of the specific ATPase inhibitor oligomycin A and of carbonyl cyanide-4(trifluoromethoxy) phenylhydrazone (FCCP) as uncoupling agent. No difference in endogenous respiration (basal) was observed between control and mutant cells, but the uncoupled oxygen consumption (FCCP) was significantly decreased in *SPART*^892dupA^ mutant cells (n=3, p=0.0306; Fig. 5A). We found that cells lacking Spartin exhibited a significant lower ATP/ADP ratio in comparison to controls, due to the concomitant decrease of ATP and increase of ADP levels (p=0.0065; Fig. 5B). We investigated the OXPHOS enzyme activities by measuring: NADH-cytochrome c oxidoreductase activity (Complex I+III); succinate dehydrogenase-cytochrome c oxidoreductase activity (Complex II+III) and NADH-DB oxidoreductase activity (Complex I). *SPART*^892dupA^ mutant cells showed a 50% decrease of Complex I+III and Complex I activities (p<0.0001 and p=0.0268 respectively; Fig. 5C, D), whereas no difference compared to controls was found for Complex II+III activity (p=0.6485; Fig. 5E). These data show that cells lacking Spartin presented a Complex I impairment. Furthermore, we observed a significant reduction (20%) in mitochondrial membrane potential in *SPART^8^*^92dupA^ cells, compared to control cells (p<0.0001; Fig. 5F) as consequence of impaired OXPHOS activity.

**Figure 5.**
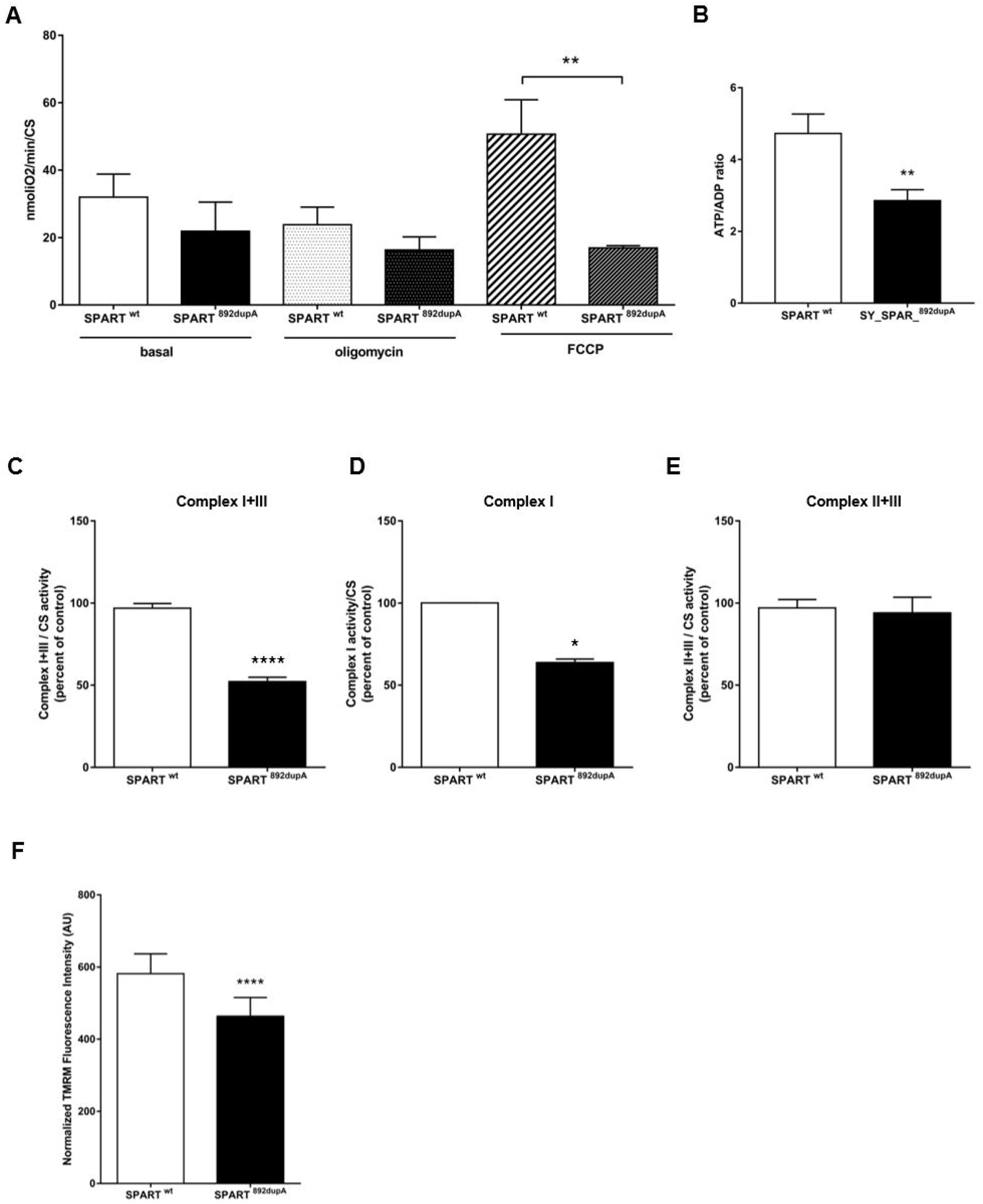
Spartin loss alters mitochondrial activity. **(A)** Oxygen consumption rates analysis in intact cells in *SPART*^wt^ (n=3 independent experiments) and *SPART*^892dupA^ cells (n=3 independent experiments). Respiration was measured in DMEM (basal respiration), in presence of oligomycin A (non-phosphorylating respiration) and in presence of FCCP (uncoupled respiration). Data were normalized on citrate synthase activity. Colour legend: white box= *SPART*^wt^ cells; black box= *SPART*^892dupA^ cells; white dotted box= *SPART*^wt^ cells oligomycin A-treated; black dotted box=*SPART*^892dupA^ cells oligomycin A-treated; white striped box= *SPART*^wt^ cells FCCP-treated cells; black striped box= *SPART*^892dupA^ cells FCCP-treated cells. **P < 0.01, means ± SEM. **(B)** ATP/ADP ratio in cellular extracts from *SPART*^wt^ and *SPART*^892dupA^ cells showing a decreased ATP/ADP ratio in mutant cells. Bars indicate standard errors. **P < 0.01, means ± SEM. **(C-E)** OXPHOS complex activity measurements. **(C)** Complex I+III activity (NADH-cytochrome c oxidoreductase activity) in cell homogenate from *SPART*^wt^ (n=3 independent experiments) and *SPART*^892dupA^ cells (n=3 independent experiments). **** P < 0.0001, means ± SEM. **(D)** Complex I activity (NADH-dehydrogenase) in cell homogenate from wild type (n=3 independent experiments) and mutant SHSY-5Y cells (n=3 independent experiments). Data were normalized on citrate synthase activity (CS). *P < 0.05, means ± SEM. **(E)** Complex II+III activity (succinate-cytochrome c oxidoreductase activity) in cell homogenate from *SPART*^wt^ (n=3 independent experiments) and *SPART*^892dupA^ cells (n=3 independent experiments). **(F)** Mitochondrial membrane potential measurement assessed with Tetramethylrhodamine (TMRM) probe. The TMRM fluorescence emission was normalized on MitoTracker Green emission. (n=12) **** P < 0.0001, means ± SEM.

### Altered pyruvate metabolism in *SPART*^892dupA^ mutant cells

Since OXPHOS respiration was impaired, we investigate a possible metabolic switch to glycolysis. We measured intracellular NADH levels and extracellular lactate. In *SPART*^892dupA^ mutated cells, the intracellular NADH level was increased in comparison to control cells (p=0.0026, Fig. 6A). HPLC analysis of extracellular culture medium did not detect any change in extracellular lactate levels (Fig. 6B). Nevertheless, in the extracellular culture medium of *SPART*^892dupA^ cells, we identified a 2.5 folds increase in pyruvate levels in comparison to controls (*SPART*^892dupA^ = 231.4±27.65 vs *SPART*^wt^ = 77.56±7.639; p=0.0241, Fig. 6C, D).

**Figure 6.**
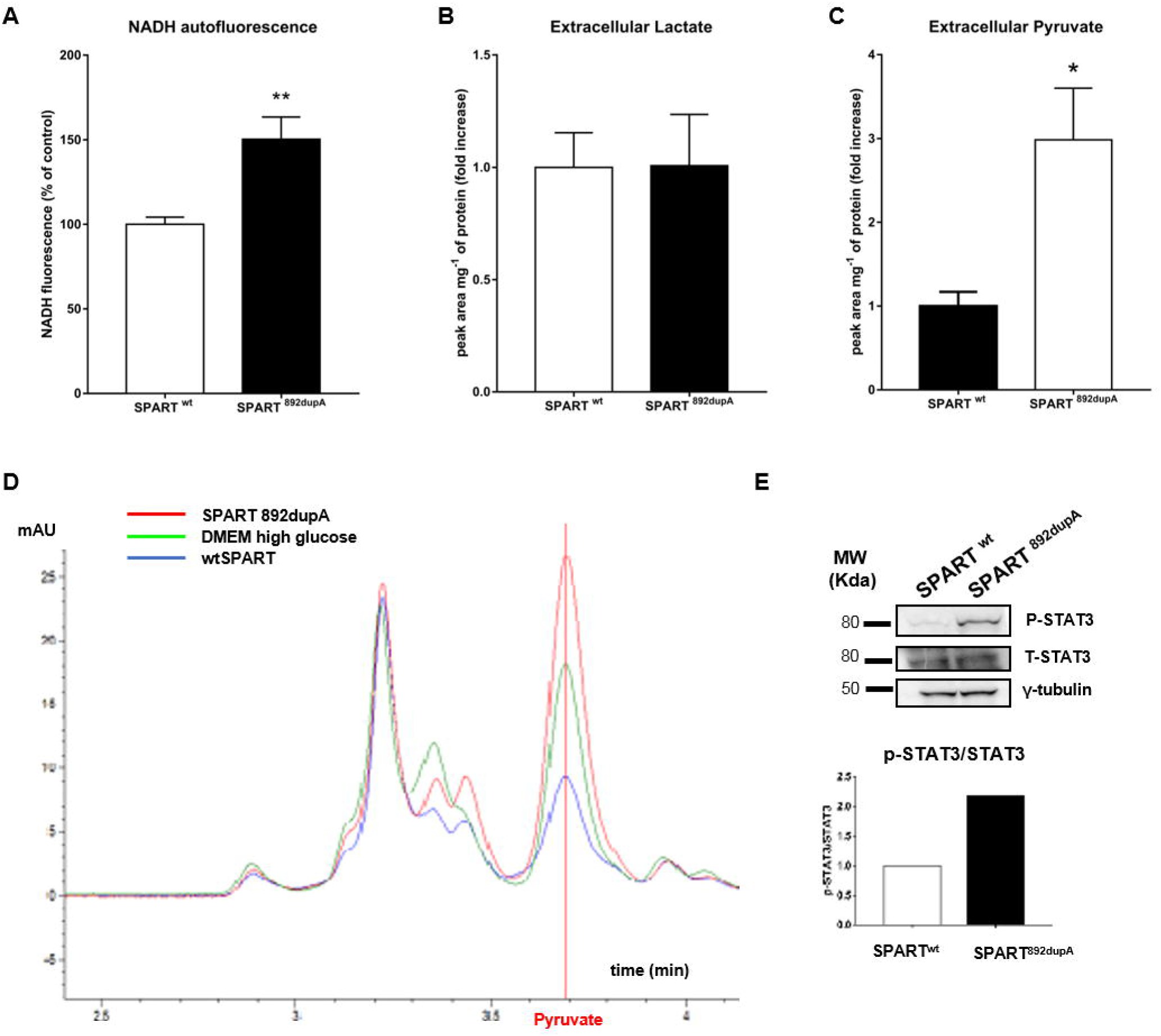
Mutant *SPART*^892dupA^ cells showed an increased NADH level and pyruvate excretion. **(A)** NADH autofluorescence measurement showing an increased level in mutant cells compared to controls. **P < 0.01, means ± SEM. **(B)** Extracellular lactate content determination by HPLC in *SPART*^wt^ and *SPART*^892dupA^ cells. The extracellular lactate content in culture cell medium was quantified after 24 hours of cell growth by HPLC analysis. The peak area corresponding to lactate was normalized on cell number. **(C)** Extracellular pyruvate production *SPART*^wt^ and *SPART*^892dupA^ cells measured by HPLC analysis after 72 hours of cell growth. The peak area corresponding to pyruvate was normalized on protein content by Bradford assay. *P < 0.05, means ± SEM. **(D)** Representative HPLC chromatograms of extracellular media from *SPART*^wt^ and *SPART*^892dupA^ cells. Red line indicates medium from *SPART*^892dupA^ cells, green line indicates medium only (DMEM high glucose) and blue line indicates medium from *SPART*^wt^ cells. Vertical red line indicates pyruvate. **(E)** Representative western blot analysis showing the expression of phosphorylated and total STAT3 in the two cell lines. Gamma tubulin was used as endogenous control. *SPART*^892dupA^ showed an increase of P-STAT3 compared to *SPART*^wt^ cells, indicating an over-activation of STAT3 in mutant cells. Graph showed the relative quantification of western blot.

### STAT3 activation in *SPART*^892dupA^ mutant cells

Decreased oxidative phosphorylation has been correlated to STAT3 activation, and constitutive activation of STAT3 in several cell models indicated a major role for this transcription factor in promoting an increase in glycolysis (Demaria *et al.*, 2010). Thus, we investigated the phosphorylation/activation status of STAT3 in mutant and control cell lines via western blot analysis for phosphorylated STAT3 and total (T-STAT3). *SPART*^892dupA^ cells showed STAT3 phosphorylation (P-STAT3), whereas no activation was observed in *SPART*^wt^ cells. Similar levels of total STAT3 were present in both cell types (Fig. 6E).

### Spartin loss increased mitochondrial Reactive Oxygen Species (ROS) and altered intracellular Ca^2+^ homeostasis

Increases in cellular superoxide production have been implicated in a variety of pathologies, including neurodegeneration (Sena and Chandel, 2012). Mitochondrial superoxide is generated as a by-product of oxidative phosphorylation and in healthy cells occurs at a controlled rate. Given the specific NADH-dehydrogenase activity (Complex I) impairment detected in mutant cells, we investigated whether ROS production was altered by Spartin loss. Intracellular ROS levels, measured with the fluorescent probe 2',7'–dichlorofluorescin diacetate (DCFDA), were significantly increased in *SPART*^892dupA^ cells (p<0.0001; Fig. 7A). Moreover, by staining live cells with MitoSOX™ Red, a highly selective probe for detection of mitochondria-specific superoxide, we found a significant increase in superoxide production in *SPART*^892dupA^ cells compared to the control (p<0.0001; Fig. 7B). To further investigate the oxidative stress status of mutated cells, we tested the expression of the major ROS-detoxifying enzymes. RT-qPCR revealed a significant reduction in expression of *CAT* (Catalase), *SOD1* (Superoxide Dismutase) and *SOD2* (mitochondrial Manganese Superoxide Dismutase) in cells lacking Spartin, compared to controls (p=0.0027 for *CAT*; p=0.0280 for *SOD1*; p=0.0172 for *SOD2*; Fig. 7C-E, respectively).

**Figure 7.**
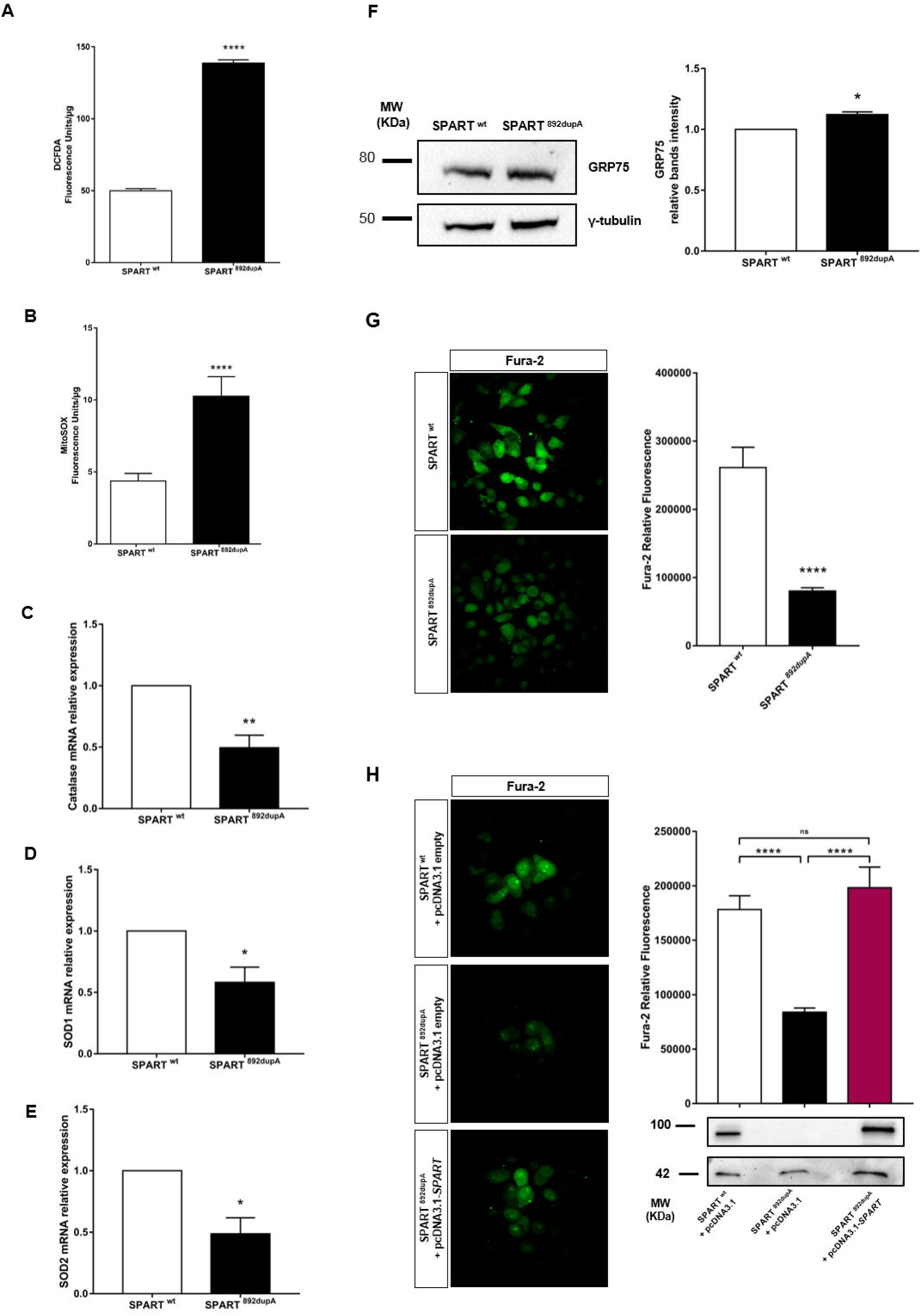
Spartin loss increases oxidative stress and alters the homeostasis of calcium. **(A)** Assessment of reactive oxygen species (ROS) production in *SPART*^wt^ (n=35) and *SPART*^892dupA^ (n=35) SH-SY5Y live cells using dichlorofluorescein diacetate (DCFDA) as fluorescent probe. **(B)** Assessment of mitochondrial superoxide production in *SPART*^wt^ (n=35) and *SPART*^892dupA^ (n=35) SHSY-5Y live cells using MitoSOX Red as specific fluorescent probe. Data were normalized on protein content using the Lowry assay. **** P < 0.0001, means ± SEM. **(C)** *CAT*, **(D)** *SOD1* and **(E)** *SOD2* mRNA relative expression in *SPART*^wt^ and *SPART*^892dupA^ cell lines. *P < 0.05, **P < 0.01, means± SEM. **(F)** Representative western blot analysis showing the expression of GRP75 protein and the relative quantification in the two cell lines. Gamma tubulin was used as endogenous control. *SPART*^892dupA^ showed a higher expression of GRP75 compared to *SPART*^wt^ cells * p < 0.05, means ± SEM. **(G)** Representative confocal microscopy images showing *SPART*^wt^ and *SPART*^892dupA^ cells stained with Fura-2-AM (right panel). Left panel showed quantification of the Ca^2+^-free Fura-2 AM relative fluorescence. *SPART*^892dupA^ (n=50) cells showed an increase of intracellular Ca^2+^ compared to control cells (n=50). **** P < 0.0001, means ± SEM. **(H)** Representative confocal microscopy images of Fura-2 AM showing *SPART*^wt^ and *SPART*^892dupA^ cells and re-expression of Spartin in mutant cells (right panel). Left panel showed Spartin re-expression evaluated by western blot and the quantification of the intracellular- free Ca^2+^ measuring the Fura-2 AM relative fluorescence in the three samples. Re-expression of Spartin rescued the intracellular Ca^2+^ concentration. **** P < 0.0001, Data are reported as the mean ± standard deviation of at least three independent experiments.

Glucose-Regulated Protein 75 (GRP75) has a major role in neuronal cells for mitochondrial function regulation and protection from stress-induced ROS (Honrath *et al.*, 2017) and physically interacts with Spartin (Milewska *et al.*, 2009). Therefore, we evaluated its protein levels, and found a higher expression of GRP75 in *SPART*^892dupA^ compared to *SPART*^wt^ cells (p=0.0327; Fig.7F).

GRP75 also coordinates the exchange and transfer of Ca^2+^, thereby affecting mitochondrial function and intracellular Ca^2+^ homeostasis (Honrath et al., 2017). In accordance, we assessed intracellular free Ca^2+^ in *SPART*^wt^ and *SPART*^892dupA^ cells by quantifying the Ca^2+^ probe Fura-2 AM-relative fluorescence. We found a significant increase of intracellular Ca^2+^ in *SPART*^892dupA^ cells, compared to *SPART*^wt^ cells (p<0.0001; Fig. 7G) consistently with previous results showing that Spartin decreased expression via transient silencing led to a dysregulation of intracellular Ca^2+^ levels (Joshi and Bakowska, 2011).

In order to provide additional evidence that these observed defects were specifically due to Spartin absence, we re-expressed Spartin in *SPART*^892dupA^ cells., Spartin re-expression rescued altered intracellular Ca^2+^ comparably to control cells (*SPART*^wt^ *vs. SPART*^892dupA^ p<0.0001; *SPART*^wt^ *vs. SPART*^892dupA+ *SPART*^ p=0.5690; *SPART*^892dupA^ *vs. SPART*^892dupA+ *SPART*^ p<0.0001; Fig. 7H, upper panel).

## Discussion

Exomic/genomic sequencing is increasingly being utilized in clinical settings for diagnostic purposes (Tan *et al.*, 2017; Krabbenborg *et al.*, 2016; Stark *et al.*, 2016).

We identified a novel homozygous insertion (c.892dupA) in the gene *SPART* detected with WES in two male sibs with syndromic short stature and developmental delay with severe speech impairment, born of a consanguineous marriage from healthy parents. This novel *SPART* mutation generated a frameshift and a premature stop codon in the Spartin protein. Loss-of-function mutations in *SPART* cause Troyer syndrome (OMIM 275900), a very rare recessive form of spastic paraplegia resulting in muscle weakness, distal atrophy, short stature and cognitive defects, due to the degeneration of the corticospinal motor neurons (Alazami *et al.*, 2015; Tawamie *et al.*, 2015; Manzini *et al.*, 2010; Patel *et al.*, 2002). Although Troyer syndrome was not firstly hypothesized, the molecular findings obtained with NGS analysis led to a thorough clinical reassessment, that identified, muscular hypotrophy in upper and lower limbs, increased muscle tone at lower limbs (distal>proximal) and brisk deep tendon reflexes. These data are in line with the findings often observed in hereditary spastic paraplegias (HSPs), where the clinical symptoms may develop later in time, therefore hindering a correct molecular and clinical diagnosis (Fink, 2013).

So far, very few *SPART* loss-of-function mutations have been reported (Bizzari *et al.*, 2017; Dardour *et al.*, 2017; Spiegel *et al.*, 2017; Butler *et al., 2016*; Alazami *et al.*, 2015; Tawamie *et al.*, 2015; Bakowska *et al.*,2008; Patel *et al.*, 2002; Supplementary Table 2). Animal (null mice) and cellular models (Spartin silencing/overexpression) indicated a role for Spartin in different processes, including neuronal survival/sprouting, cell division and mitochondrial stability (Eastman *et al.*, 2009; Edwards *et al.*, 2009; Bakowska *et al.*, 2007; Bakowska *et al.*, 2005; Ciccarelli *et al*., 2003). Perturbations of the mitochondrial dynamics or energetic failures are pathological mechanisms underpinning many neurodegenerative disorders, often with overlapping clinical features such as Hereditary Spastic Paraplegia (HSP), Alzheimer’s disease, Parkinson’s disease, Amyotrophic Lateral Sclerosis (ALS) and Huntington’s disease (Johri and Beal, 2012). Neurons, especially those with long axons, such as peripheral sensory neurons and motor neurons, are more susceptible to neurodegeneration, since they are more sensitive to alterations in mitochondrial network (Su *et al.*, 2010). All neuronal cellular processes are energy demanding and require significant active mitochondria, with a well-regulated balance between motile and stationary mitochondria.

On the basis of this evidence, we focused our functional studies to investigate the effect of *SPART* c.892dupA on mitochondrial network integrity and mitochondrial activity, along with survival and differentiation. We used two different cell models: hNSCs (silenced for *SPART*) and SH-SY5Y cell line genome-edited via CRISPR/Cas9 technology to introduce the mutation c.892dupA.

We found that both hNSCs silenced for *SPART* and SH-SY5Y cells carrying the *SPART* loss-of-function mutation presented increased cell differentiation, with impaired neuronal growth/survival, and significant neurite outgrowth/length (Fig. 2), in line with the data observed in animal models (Renvoisé *et al.*, 2012). Gene expression analysis in zebrafish embryos identified high expression of the *SPART* homologous (*spg20b*) through the initial phases, in particular during cleavage and blastula periods - 0.75-5 hours post fertilization (Supplementary Fig. 3A). In adult zebrafish brain, *spg20b* expression was 18 times higher than in heart and liver tissues (Supplementary Fig. 3B).

We observed an altered mitochondrial network in the absence of Spartin (Fig.4). Specifically, we observed a decrease in the number of mitochondria present in neurite-like extensions of mutant cells compared to control cells, which might exert a detrimental effect on dendrites and axons during synaptic transmission (Court *et al.*, 2012; Cheng *et al.*, 2012).

In the mutant cells we identified a specific mitochondrial impaired respiration with impaired ATP synthesis due to a decreased Complex I activity (Fig.5), possibly contributing to the observed reduced mitochondrial membrane potential and increased oxidative stress (with concomitant decreased expression of ROS detoxifying enzymes; Fig.7). These data demonstrate that Spartin depletion leads to mitochondrial Complex I deficiency, contributing to neurodegeneration by ROS production (Fig. 8A, B).

**Figure 8.**
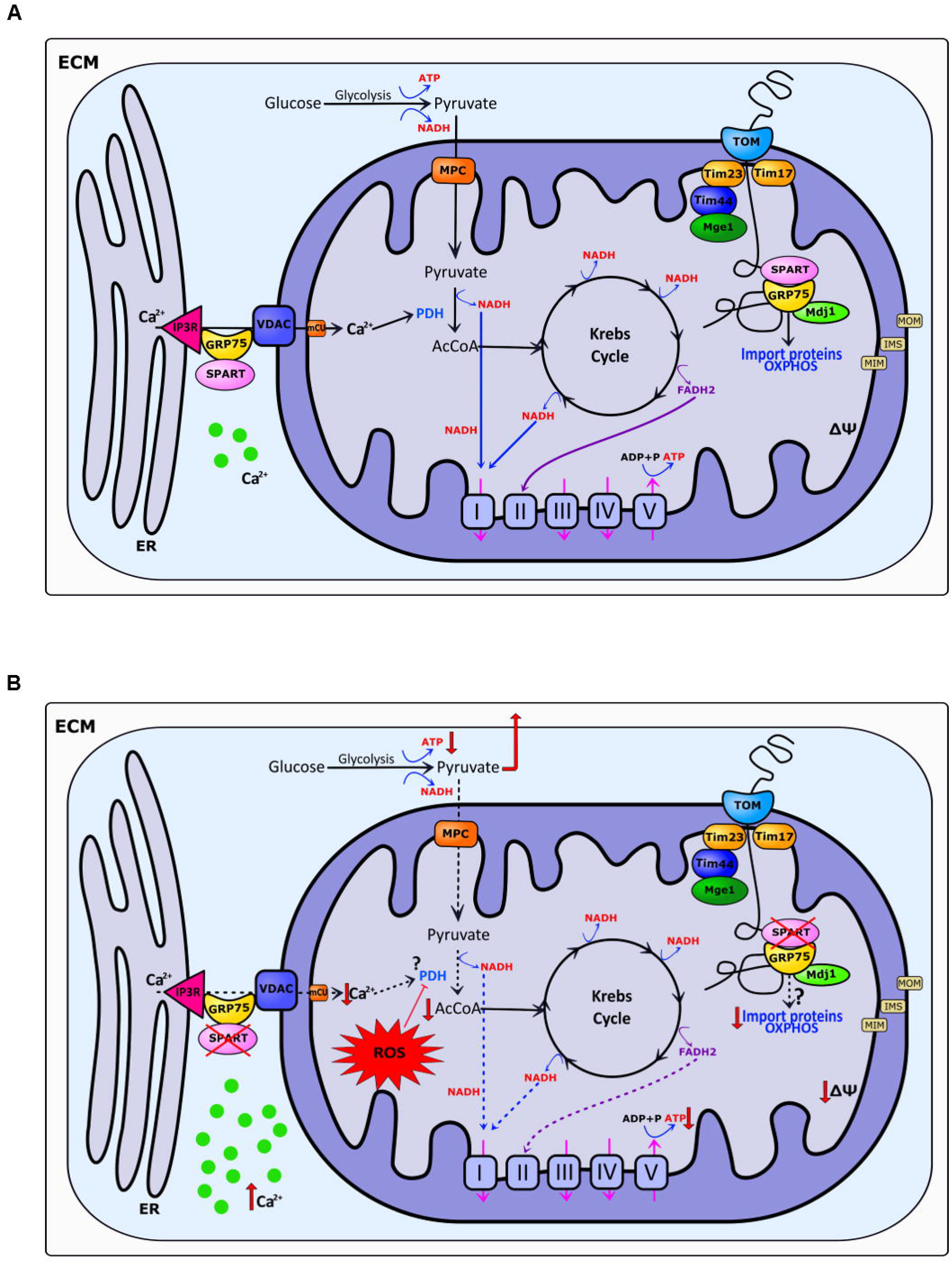
Model of Spartin functions in mitochondrial metabolism. **(A)** Spartin interacts with GRP75, modulating the import of mitochondrial proteins encoded by the nucleus via the TIM-TOM complexes into the mitochondria (Milewska *et al*., 2009), allowing a normal ATP production through Krebs’s cycle and OXPHOS activity (Complex I-V). **(B)** Spartin loss, as observed in our experimental settings, determines an energetic failure due to an altered Complex I activity, with a decreased ATP production, and a halt in mitochondrial oxidative phosphorylation leading to decreased mitochondrial membrane potential, and increased oxidative stress, due to enhanced production of mitochondrial ROS. Increased oxidative stress/increased ROS are known inhibitors of PDH activity (Gray *et al.*, 2014). Moreover, Spartin loss results in a reduction of mitochondrial Ca^2+^ levels (Joshi and Bakowska, 2011) which could alter the activity of PDH. Accordingly, we observed that pyruvate, which enter in mitochondria through the mitochondrial pyruvate carrier (MPC), was not efficiently converted into acetyl-coA (AcCoA), Krebs’s cycle substrate, thus accumulating and excreted from *SPART^892dupA^* mutant cells. Therefore, we hypothesize that the energetic failure observed in absence of Spartin, probably related to an impaired mitochondrial protein import of the nuclear-encoded subunits, is pivotal in generating the neurodegenerative defects observed in neurons in Troyer syndrome Abbreviations: MPC, mitochondrial pyruvate carrier; PDH, pyruvate dehydrogenase; Δψ, mitochondrial membrane potential; MOM, mitochondrial outer membrane; MIM, mitochondrial inner membrane; IMS, inter-membrane space; ECM, extracellular matrix. I-V: mitochondrial OXPHOS complexes; TOM, translocase of the outer membrane; IP3R, Inositol trisphosphate receptor, Phosphatidylinositol 3-phosphate; VDAC, Voltage-dependent anion-selective channel.

Another form of HSP, due to mutations in the spastic-paraplegia-7 gene (SPG7), encoding for Paraplegin, arises because of mitochondrial Complex I impairments (Casari *et al.*, 1998). Paraplegin forms large complexes in the inner membrane of mitochondria (for a review see Patron *et al*., 2018). Loss-of-function mutations in *SPG7* gene lead to a defective assembly of Complex I and consequent defective respiratory chain activity (Casari *et al.*, 1998). As demonstrated for Paraplegin, Spartin interacts with GRP75 (Fig. 8A), member of mitochondrial complex for the import of nuclear-encoded proteins into the mitochondria (Milewska *et al.*, 2009)z. Consistently with previous reports for SPG7 (Atorino *et al.*, 2003)., we can hypothesize a role for Spartin similar to Paraplegin explaining the observed Complex I deficit with assembly or stability defects consistently with this hypothesis the increased levels of GRP75 in *SPART* mutant cells (Figure 7F), suggest a compensatory effect for its absence.

We identified for the first time an excess of pyruvate in the context of Troyer syndrome, since. we found an elevated release of pyruvate in the medium from *SPART*^892dupA^ cells (Fig. 6C). Pyruvate is the end-product of glycolysis which independently from its cellular source, is directed into mitochondria through mitochondrial pyruvate carrier (MPC) located in the mitochondrial inner membrane (MIM). Here, pyruvate functions as fuel input for the citric acid cycle and for mitochondrial ATP generation. Disruption in pyruvate metabolism affects tissues with high demand for ATP. The nervous system is particularly vulnerable because of its high demand of carbohydrate metabolism for ATP generation (Gray *et al.*, 2014). Our data suggest that increased pyruvate excretion underlies an increased glycolytic pathway, with a subsequent halt in the carbon flux due to impaired Complex I activity, increased ROS production and damage to the cells (Fig. 8A, B). We also detected a constitutive phosphorylation (hence activation) of STAT3 associated to Spartin loss (Fig. 6E). Constitutive STAT3 activation has been shown to promoted increase in glycolysis and decrease in oxidative phosphorylation (Demaria *et al.*, 2010) with a role in faster neurite outgrowth (Zhou and Too, 2011). STAT3 contrasts the activity of pyruvate dehydrogenase (PDH), a key enzyme of glycolysis that converts pyruvate into acetylCoenzyme A (AcCoA) (Demaria *et al*., 2010). Conversely, Spartin has been shown to promote PDH activity in yeast (Ring *et al.*, 2017).

Our data are consistent with an effect of Spartin loss on pyruvate metabolism by dysregulation of PDH activity (Fig. 8A, B). PDH inhibition was also reported in presence of oxidative stress (Tabatabaie *et al*., 1996; Hurd *et al*., 2012; Liemburg-Apers *et al*., 2015) and in decreased mitochondrial Ca^2+^ levels (Holness and Sugden, 2003). It is worth noting that the GRP75/VDAC (Voltage-dependent anion channel) complex regulates mitochondrial Ca^2+^ intake from the endoplasmic reticulum thereby affecting mitochondrial function and intracellular Ca^2+^ homeostasis (Szabadkai *et al.*, 2006; Honrath et al., 2017). Consistently, we observed an increase in intracellular Ca^2+^ in SPART^892dupA^ cells compared to SPART^wt^ cells (Fig. 7G, which was restored to normal levels with the re-expression of Spartin (Fig. 7H). In accordance to Joshi and Bakowska (2011), these results suggest that Spartin loss itself affects trafficking and buffering cytosolic and mitochondrial Ca^2+^ via GRP75 interaction.

In summary, we showed for the first time that Spartin depletion causes a decreased activity of mitochondrial Complex I coupled to increased pyruvate excretion. Our data suggest that Spartin coupled with GRP75 might modulate mitochondrial protein import and Ca^2+^ levels, maintaining low levels of ROS and a normal ATP production (Fig. 8A). Spartin loss determines an energetic failure, due to altered Complex I function, with increased ROS production possibly leading to a reduction of PDH activity (Fig. 8B; Gray *et al.*, 2014). Therefore, pyruvate is not efficiently converted into acetyl-coA (AcCoA) and is accumulated and excreted from *SPART* mutant cells (Fig. 8B).

We suggest that Troyer syndrome to be a mitochondrial disease. We propose that the observed neuronal phenotypes result from defective protein assembly within mitochondria or defective mitochondrial Complex I function, coupled to an excess of pyruvate and ROS production. Our results are relevant to interpret the molecular mechanism of other neurological syndromes/disorders associated to Complex I impairments, such as Spastic paraplegia 7.

As the expansion of personalized medicine proceeds, with increasing potential for active analyses of genomic data, the early identification of molecular and genetic defects can lead to a better clinical refinement and the possibility of applying timely targeted therapies. Most importantly, it is crucial to carry out the functional characterization of proteins and mutations, which is the only way to move forward translational work from gene identification.

## Supporting information

## Acknowledgements

We thank all patients that participated in the study. We thank Dr. A. Astolfi, Ms. S. Di Battista and Ms. M. Cratere for technical help in sequencing analysis and cell culture, Ms. A. Martinelli, and Dr. M. Gostic for help in zebrafish analysis.

## URL

Exome Aggregation database (ExAc): http://exac.broadinstitute.org/

Genome Aggregation database (gnomAD): http://gnomad.broadinstitute.org/

Eurexpress: http://www.eurexpress.org

Human Protein Atlas: https://www.proteinatlas.org/ENSG00000133104-SPG20

CRISPR design: crispr.mit.edu

## Abbreviation

AcCoA: Acetyl-coA
ALS: Amyotrophic Lateral Sclerosis
BMP: Bone Morphogenic Protein
CAT: Catalase
Complex I: NADH dehydrogenase
Complex I+III: NADH-cytochrome c reductase activity
Complex II+III: succinate-cytochrome c reductase activity
CS: Citrate Synthase
DAPI: 4′,6-diamidino-2-phenylindole
DB: Decylbenzoquinone
DCFDA: 2′,7′–dichlorofluorescin diacetate
DDS: Deciphering Consortium
DMEM: Dulbecco’s modified Eagle’s medium
DTNB: 5,5′-dithiobis-2-nitrobenzoic acid
ECM: Extra-Cellular Matrix
ExAc: Exome Aggregation database
FCCP: carbonyl cyanide-4-(trifluoromethoxy) phenylhydrazone
GH: Growth Hormone
gnomAD: Genome Aggregation database
gRNA: guide RNA
GRP75: Glucose-Regulated Protein 75
hESC: human embryonic stem cell
hNSC: human neural stem cell
hpf: hours post fertilization
HPLC: High Performance Liquid Chromatography
HSP: Hereditary Spastic Paraplegia
IMS: Inter-Membrane Space
IUGR: Intra-Uterine Growth Restriction
MIM: Mitochondrial Inner Membrane
MIT: Microtubule Interacting and Trafficking
MOM: Mitochondrial Outer Membrane
MPC: Mitochondrial Pyruvate Carrier
MTG: Mitotracker Green
mΔψ: Mitochondrial transmembrane potential
OXPHOS: oxidative phosphorylation
PDH: Pyruvate Dehydrogenase
pSpCas9n(BB)-2A-GFP: Cas9 from S. pyogenes with 2A-EGFP
pSpgRNA: S. pyogenes Cas9 guide RNA
ROH: Runs Of Homozygosity
ROS: reactive oxygen species
SD: standard deviations
SDS-PAGE: sodium dodecyl sulfate polyacrylamide electrophoresis
SEM: Standard Error of Mean
SNPs: Single Nucleotide Polymorphisms
SOD1: Superoxide Dismutase
SOD2: Manganese Superoxide Dismutase
ssODN: single stranded-oligonucleotide
TBS: Tris Buffered Saline
TMRM: Tetramethylrhodamine methyl ester
UPD7: Uniparental disomy 7
WES: Whole Exome Sequencing

## Materials and Methods

### High-throughput SNP genotyping and Whole Exome Sequencing (WES)

#### High-Throughput SNP genotyping

High-throughput SNP (Single Nucleotide Polymorphisms) genotyping has been performed on Illumina Infinium HD Assay Gemini platform (Illumina, San Diego, CA, USA), according to manufacturer’s protocol, starting from 400 ng of genomic DNA form peripheral blood. Genotypes were converted into PLINK format with custom scripts. PLINK v1.07 (http://ngu.mgh.harvard.edu/~purcell/plink/) was used to isolate individual Runs Of Homozygosity (ROH) that showed > 1 Mb overlap Mb overlap between the three affected siblings (Bonora *et al*, 2015).

#### Whole Exome Sequencing (WES)

WES was performed on genomic DNA extracted from peripheral blood (QIAGEN mini kit) from the two affected brothers. Genomic DNA libraries, starting from 100 ng genomic DNA, were prepared using the Illumina Pair-End Nextera Kit (Illumina, San Diego, CA USA) and library was enriched for exomic sequences using the Nextera coding exome kit. The captured regions were sequenced on the Illumina HiScanSQ platform for 200 cycles (100 cycles paired-ends; Illumina). The read files were aligned to hg19 version of the human genome sequencing, annotation and variant prioritization was performed according to our internal pipeline for exome annotation as previously reported (Graziano *et al.*, 2015). The identified variants were confirmed by PCR and direct sequencing.

### Cell lines

SH-SY5Y cells (ATCC, UK) were cultured in Dulbecco’s modified Eagle’s medium (DMEM, Euroclone, Milan, Italy) supplemented with 10% (v/v) fetal bovine serum, 100 U/mL penicillin and 100 µg/mL streptomycin (supplements were purchased from Sigma-Aldrich, St. Louis, MO USA). Human neural stem cells (hNSC), derived from the NIH approved H9 (WA09) human embryonic stem cells (hESCs), were grown in 6-well plates coated with CTS™ CELLstart™ Substrate (Gibco) and maintained in KnockOut D-MEM/F-12 with 2mM of GlutaMAX-I supplement, 20 ng/ml of bFGF, 20 ng/ml of EFG and 2^%^ of StemPro^®^ Neural Supplement (all reagents were purchased from Thermo Fisher Scientific, Waltham, MA USA). All cells were grown in a humidified incubator with 95% air and 5% CO_2_ at 37°C.

### Silencing SPART in Neural Stem Cells (hNSC)

To transiently knock-down *SPART*, hNSC were transfected every 36 hours with a combination of 3 siRNAs (Thermo Fisher Scientific; see Supplementary Table 1 for the sequences), using Lipofectamine3000 (Thermo Fisher Scientific) according to the manufacturer’s instruction. At day 0, 4 and 8 cells were collected and processed for western blot and imaging analysis.

### Generation of SPART c.892dupA knock-in SH-SY5Y cell line

The *SPART* mutation was generated in SH-SY5Y genome using, guide RNAs (gRNAs) designed with the CRISPR Design Tool (mit.edu.crispr) (Hsu *et al.*, 2013). The two guides were cloned into the pSpgRNA (S. pyogenes Cas9/dCas9 guide RNA) expression vector (#47108; Addgene, Cambridge, MA USA; Perez-Pinera *et al.*, 2013) and sequenced. Sequences of gRNAs and single stranded-oligonucleotide (ssODN) carrying the variant c.892dupA are reported in Supplementary Table 1. Cells were plated in a T25 flask and when 80% confluent, they were transfected with 4.7 µg of pSpCas9n(BB)-2A-GFP (PX458, #48140; Addgene; Ran et al., 2013), 0.8 µg of each gRNA expression plasmid and 10uM of ssODN with Lipofectamine^®^3000 (Thermo Fisher Scientific) according to the manufacturer’s instruction. After 28 hours, cells were sorted with an automated Fluorescence-activated cell sorting (FACS) (Influx, Becton Dickinson) and single cells plated in 96-wells plates coated with Poly-D-Lysine (Sigma). Clones were amplified and screened by PCR and direct sequencing of the target region. A clone carrying the specific change and with no off-target mutations was selected for the analysis (hence defined *SPART*^892dupA^). The SH-SY5Y clone that underwent the same CRISPR/Cas9 genome editing approach but did not carry any change was used as control cell line (hence defined *SPART*^wt^ throughout the text).

### Western blotting

Cells were lysed in ice-cold RIPA buffer: 50 mM HEPES (EuroClone), 1 mM EDTA (Sigma-Aldrich), 10% glycerol (Fisher Fisher Scientific), 1% Triton X-100 (Sigma-Aldrich), 150 mM NaCl in the presence of proteases and phosphatases inhibitors (Sigma-Aldrich). Total protein was measured using the Lowry protein assay kit (Biorad DC Protein Assay; Biorad, Hercules, CA USA) according to the manufacturer’s instruction. Protein samples (70 μg) were subsequently separated on 10% sodium dodecyl sulfate polyacrylamide electrophoresis (SDS-PAGE) gels or on 4-20% pre- cast SDS-PAGE gels (Biorad). Gels were then electro-transferred onto nitrocellulose membranes (Trans-Blot Turbo Transfer System, Biorad). Membranes were blocked in Tris Buffered Saline (TBS) with 1% Casein (Biorad) for 1 hour at room temperature and incubated with primary antibodies at 4°C overnight. Membranes were washed three times in Tris-buffered saline containing 0.1% Tween and incubated with peroxidase-conjugated secondary antibodies for 45 minutes at room temperature. Bands were visualized using WESTAR Supernova (Cyanagen, Bologna, Italy) and detected with the ChemiDoc™ XRS+ (Biorad). Densitometric analysis was performed with ImageLab software (Biorad). Primary antibodies used were: GAPDH (mouse, 1:10,000; Abcam, Cambridge, UK), γ-tubulin (mouse, 1:10,000; Sigma-Aldrich), T-STAT3 (mouse, 1:500; OriGene, Rockville, MD USA), P-STAT3 (rabbit, 1:500; Cell Signaling, Leiden, Netherlands), Spartin (rabbit, 1:1,000; ProteinTech, Rosemont, IL USA) and GRP75 (goat, 1:500; Santa Cruz, CA USA). Peroxidase-conjugated secondary antibodies used were anti-mouse IgG (1:5,000), anti-rabbit IgG (1:5,000) from Sigma-Aldrich), and anti-goat IgG (1:5,000; Dako, Glostrup, Denmark).

### Immunofluorescence microscopy

Cells were plated in µ-Slide 8 Well (Ibidi) coated with Poly-D-Lysine (Sigma-Aldrich). When 80% confluents, were fixed in 4% paraformaldehyde in PBS for 10 minutes at 4°C. Samples were blocked and permeabilized in 10% newborn calf serum (Sigma-Aldrich), 0.3% Triton X-100 (Sigma-Aldrich) in PBS for 1h at room temperature. Samples were incubated 16 hours at 4°C in primary antibody diluted in 5% newborn calf serum, 0.15% Triton X-100 in PBS. After three 30-minutes washes in PBS, sample were incubated 16 hours at 4°C in secondary antibodies diluted in 5% newborn calf serum, 0.15% Triton X-100 in PBS. After three 30-minutes washes in PBS, samples were mounted in Fluoroshield with DAPI (4′,6-diamidino-2-phenylindole; Sigma-Aldrich). Microscopies used for imaging were Leica and Axiovert 200 inverted microscope (Carl Zeiss, Oberkochen, Germany). Primary antibodies were used against: β-III tubulin (mouse, 1:500; Abcam), Nestin (mouse, 1:300; Abcam) and PGP9.5 (rabbit, 1:300; Thermo Fisher Scientific). Alexa Fluor-conjugated secondary antibodies used were: Alexa Fluor 488 goat anti-rabbit IgG, Alexa Fluor 488 donkey anti-mouse, Alexa Fluor 555 goat anti-rabbit and Alexa Fluor 555 donkey anti-mouse (all diluted at 1:800; Abcam).

### RNA isolation and quantitative PCR (qPCR) in cell lines

Total RNA was isolated from SH-SY5Y cultures using the RNeasy Mini Kit (Qiagen, Hilden, Germany). cDNA from 1µg of *SPART*^wt^ and *SPART*^892dupA^. RNA was synthesized using the SuperScript™ VILO™ cDNA Synthesis Kit (Thermo Fisher Scientific). Quantitative PCR was performed using SYBR^®^ Green master mix (Biorad). All samples were run in triplicate on the ABI7500 Fast PCR machine (Thermo Fisher Scientific). Melting curve analysis for each primer pair was carried out to ensure specific amplification. Relative mRNA expression levels of *CAT, SOD1, SOD2,* were normalized to the house-keeping gene β-actin using the ΔΔCt method. Primers are reported in Supplementary Table 1.

### Quantification of neurite outgrowth and elongation

Cells were seeded on poly-L-lysine (Sigma-Aldrich) coated glasses and immunofluorescence of the neuronal marker PGP9.5 have been performed as described above. Mean neurite number and length was measured using the NeuronGrowth plugin (ImageJ, National Institute of Health, Bethesda) by tracing the individual neurites of cells randomly analyzed. For each experiment twenty cells have been examined.

### Spheroid formation assay

We generated a matrix-free SH-SY5Y spheroid cultures by seeding 2×10^4^ *SPART*^wt^ and *SPART*^892dupA^ SH-SY5Y in ultra-low attachment µ-slide 8-wells (Ibidi, Munich, Germany). Cells were grown to allow the spontaneous formation of spheroids. After 4 days, spheroids were examined and photographed using a phase contrast microscope and then RNA was extracted. Spheroids morphology was measured using ImageJ software.

### ROS quantification

#### Intracellular ROS measurement using DCFDA

Control and *SPART*^892dupA^ SH-SY5Y cell lines were seeded at 5 x 10^3^ cells/well and incubated overnight. Cells were treated with 10 µM DCFDA (2',7'–dichlorofluorescin diacetate) dissolved in medium for 1 hour. Finally, cells were washed with PBS and the fluorescence emission from each well was measured (λexc = 485 nm; λem = 535 nm) with a multi-plate reader (Enspire, Perkin Elmer, Waltham, MA USA) and normalized on protein content by Lowry assay. Data are reported as the mean ± standard deviation of at least three independent experiments.

#### Mitochondrial ROS measurement using MitoSOX

Mitochondrial superoxide production was measured using MitoSOX™ Red (Molecular Probes, Thermo Fisher Scientific) following manufacturer instructions with minor modifications. Briefly, cells were seeded in 96-well plates (OptiPlate black, Perkin Elmer) at 5 x 10^3^ cells/well and incubated overnight to allow adhesion. After this time, cells were treated with 5 µM MitoSOX Red dissolved medium for 30 minutes. Cells were then washed twice times with warm PBS and the fluorescence emission from each well was measured (λexc=510 nm; λem=580 nm) with a multi-plate reader (Enspire, PerkinElmer) and normalized on protein content by Lowry assay. Data are reported as the mean ± standard deviation of at least six independent experiments.

### Mitochondrial network and morphology assessment via live cell imaging

To visualize the mitochondrial network in live cells, 3×10^4^ cells were plated in µ-Slide 15 Well (Ibidi, Germany) coated with Poly-D-Lysine (Sigma-Aldrich) in 50 μL of complete medium and incubated at 37 °C in a humidified atmosphere with 5% CO2. After 24 h, cells were transfected with a GFP protein (CellLight™ Mitochondria-GFP, BacMam 2.0, ThermoFisher Scientific) targeting mitochondria following manufacturer’s instructions. Mitochondrial network morphology was assessed by live cell imaging, using a Nikon C1si confocal microscope (Nikon, Tokio, Japan) following Dagda et al., 2009 with minor modification using NIH ImageJ software. Briefly, the green channel was subjected to both a background subtraction with a radius of 10 pixels and a median filter to reduce noise. An automatic threshold value was then applied to isolate the particles. The mean area/perimeter ratio was employed as an index of mitochondrial interconnectivity.

### Mitochondrial oxygen consumption

To measure mitochondrial oxygen consumption in *SPART*^wt^ and *SPART*^892dupA^ cells, 1.5×10^6^ for each cell line were harvested at 70-80% confluence, washed in PBS, re-suspended in complete medium and assayed for oxygen consumption at 37°C using a thermostatically regulated oxygraph chamber (Instech Mod.203, Plymouth Meeting, PA, USA) in 1.5 ml of culture medium. Basal respiration was compared with the one obtained after injection of oligomycin (1 μM) and FCCP (1– 6 μM). Antimycin A (5 µM) was added at the end of experiments to completely block the mitochondrial respiration. The respiratory rates were expressed in μmol O_2_/min/mg of protein referring to 250 nmol O_2_/ml of buffer as 100 % at 30°C (Estabrook, 1967). Data were normalized to protein content determined by the Lowry assay.

### ATP and ADP synthesis determination

Nucleotides were extracted and detected following Jones DP. 1981, using a Kinetex C18 column (250 × 4.6 mm, 100 Å, 5 μm; Phenomenex, CA). Absorbance (260 nm) was monitored with a photodiode array detector (Agilent 1100 series system). Nucleotide peaks were identified by comparison and coelution with standards, and quantification by peak area measurement compared with standard curves.

### Respiratory chain complex activities

Cell lysate was resuspended in a 20 mM hypotonic potassium phosphate buffer (pH 7.5) followed by spectrophotometric analysis of mitochondrial complexes activity at 37°C using a Jasco-V550 spectrophotometer equipped with a stirring device. Complex I activity was measured in 50mM phosphate buffer at 340nm (ε=6.22 mM ^−1^ cm^−1^) after the addition of 150µg of cell lysate, 1mM KCN, 10µM antimycin a, 2.5mg fatty acid-free BSA, 100µM NADH, 60µM decylbenzoquinone (DB). Complex I (NADH dehydrogenase) specific activity was obtained by inhibiting complex I with 10µM rotenone. The succinate-cytochrome c reductase activity (II+III activity) was measured in 50mM phosphate buffer at 550 nm (ε=18.5 mM ^−1^ cm^−1^) after the addition of 100µg of cell lysate, 1mM KCN, 20mM succinate and 50µM oxidized cytochrome c. The specific complex II+III activity was obtained by inhibiting complex II with 500µM TTFA. The NADH-cytochrome c reductase activity (I+III activity) was measured in 50mM phosphate buffer at 550 nm (ε=18.5 mM^−1^ cm^−1^) after the addition of 100µg of cell lysate, 1mg/ml of fatty acid-free BSA, 1mM KCN, 50µM cytochrome c and 200µM NADH. The specific complex I+III activity was obtained by inhibiting complex I with 10µM rotenone. Citrate synthase activity was measured in 100mM TRIS buffer with 0,1% Triton X-100 at 412 nm (ε=13,600 M^−1^cm^−1^) after the addition of 30µg of cell lysate, 0.1 mM acetyl-CoA, 0.5 mM oxaloacetate, and 0.1 mM 5,5′-dithiobis-2-nitrobenzoic acid (DTNB).

### Mitochondrial transmembrane potential *(mΔψ)*

Mitochondrial transmembrane potential and mass were measured following Kirk *et al*., 2014 with minor modifications. Briefly, Cells were seeded at the density of 10000 cell/well in 96-well culture plates (OptiPlate Black, Perkin Elmer). After 24 hours, cells were loaded with 50 nM tetramethylrhodamine methyl ester (TMRM, 544ex; 590em, Thermo Fisher) and 25 nM MitoTracker Green (MTG, 490ex; 516em, Thermo Fisher) for 30 minutes and washed twice with PBS. The fluorescence emission from each well was measured with a multi-plate reader (Enspire, PerkinElmer). TMRM fluorescence emission intensity was normalized on MitoTracker Green fluorescence.

### NADH quantification

NADH autofluorescence measurements was performed as described by Frezza and colleagues (Frezza *et al.*, 2011) with minor modifications. Briefly, cells were seeded at a density of 3×10^3^ cells per well in 15 wells µ-Slides (Ibidi, Germany) following manufacturer’s instructions and incubated overnight to allow adhesion. Images were collected with a Nikon C1si confocal microscope equipped with UV laser. NADH quantification was performed using ImageJ software after background subtraction.

### Lactate and pyruvate quantification

Extracellular lactate was determined by HPLC (High Performance Liquid Chromatography). Briefly, cells were seeded in 6 well dishes and after 72 hours the culture medium was collected for HPLC analysis. Prior to injection, the culture medium was diluted 1:10 in mobile phase and centrifuged at 14000 g for 5 minutes at 4°C. The supernatant was then injected manually into the HPLC system. The separation of the different metabolites was achieved using a C18 column (Agilent ZORBAX SB-Phenyl, 5 µm, 250×4.6 mm), using a mobile phase consisting of 50 mM KH_2_PO_4_, pH 2.9, at a flow rate of 0.8 ml/min. Lactate and pyruvate were detected using an Agilent UV detector set to 210 nm and quantified using Agilent ChemStation software. The retention time was determined by injecting standard solution. All injections were performed in triplicate. The peak area was normalized on protein content measured by Bradford assay.

### Measurement of intracellular Ca2+ in live cells using Fura-2

Intracellular calcium level was assessed in live cells using Fura-2 AM probe (Thermo Fisher) following manufacturer instruction. The emission of the calcium-free probe was measured using a Nikon C1si confocal microscope (Nikon, Tokio, Japan) at λ 380 nm excitation and λ 515nm emission. The quantification of fluorescence intensity was carried out using the ImageJ software (NIH), where at least 50 cells per condition were analyzed.

### Rescue of the phenotype with Spartin wild-type

Human *SPART* coding sequence was PCR-amplified from SH-SY5Y genomic DNA using the KAPA HiFi HotStart Taq Polymerase (Kapa Biosystems, Roche Diagnostic; Mannheim, Germany) according to the manufacturer's instructions. Primers are reported in Supplementary Table 1. The amplified fragment was digested with *XhoI* and *HindIII* (New England Biolabs, Hitchin, UK), cloned in pcDNA3.1 vector and sequenced to verify the correct insertion. For rescue experiments, 4×10^5^ *SPART*^wt^ and *SPART*^892dupA^ cells were plated. The plasmids expressing wild-type *SPART* (3 μg/experiment) was transfected into *SPART*^892dupA^ cells using Lipofectamine 3000 (Life Technologies) following the manufacturer’s instructions. In parallel, the pcDNA3.1 empty vector (3 μg/experiment) was transfected into *SPART*^wt^ and *SPART*^892dupA^ cells. Forty eight hours after transfection, cells were pelleted and washed twice with PBS. Western blot analysis to verify Spartin overexpression, determination of ATP and ADP synthesis and assessment of intracellular Ca^2+^ were performed as previously described.

### Gene expression of spg20b in zebrafish developmental stages and zebrafish adult tissues

Total RNA from developmental stages between 16–32 cells, up to 120 hours post-fertilization (hpf) was extracted using the RNeasy Mini kit according to the manufacturer’s instructions (QIAGEN) using at least 50 embryos at each stage. Heart, liver and brain were dissected from 5 adult fish, flash-frozen on dry ice and stored at -80°C until the RNA was extracted. Animals were handled following the guidelines from European Directive 2010/63/EU and euthanised with Schedule 1 procedures of the Home Office Animals (Scientific Procedures) Act 1986. RNA was synthesized using the SuperScript™ VILO™ cDNA Synthesis Kit (Thermo Fisher Scientific) following the manufacturer protocol. Gene expression level of *spg20b* was assessed by quantitative PCR (qPCR) conducted with the SYBR Green master mix (BioRad). All samples were run in triplicate on the ABI7500 Fast PCR machine (Thermo Fisher Scientific). Relative mRNA expression level of *spg20b* was normalized to the eukaryotic translation eef1a1l2 as endogenous control gene. Primers are reported in Supplementary Table S1. All zebrafish studies were approved by the Animal Welfare and Ethics Committee at the University of St. Andrews.

### Images analysis

Images were analyzed with ImageJ (public software distributed by the National Institutes of Health) and Chemidoc (Biorad).

### Statistical analysis

Statistical analysis was conducted with Prism 7 (GraphPad, San Diego, CA, USA). All experiments were carried out at least in triplicates. Results are expressed as the mean ± SEM. Unpaired Student’s t-test was used to determine the differences between groups. Data analysis of the rescue experiments was performed using ANOVA test, with Tukey-multiple comparison test. A p-value <0.05 (two-tailed) was considered statistically significant.

---

**Supplementary Figure 1.**
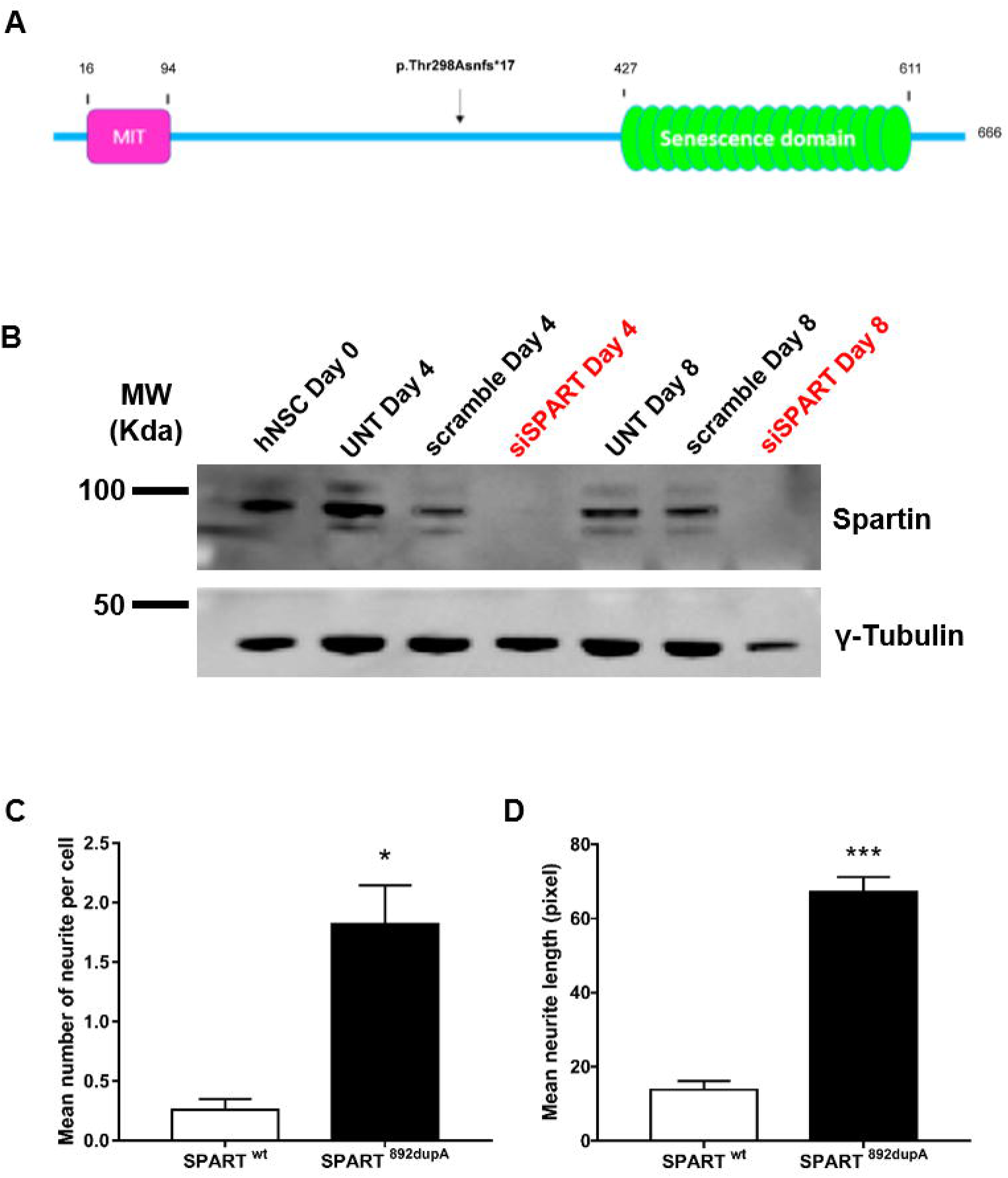
Spartin protein analysis in neural cell models. **(A)** Representative image of Spartin protein showing the Microtubule Interacting and Trafficking domain (MIT) (aa 16-94) and the senescence domain (aa 427-611). The mutation identified in the two brothers is indicated by an arrow (duplication of nucleotide 892 at amino acid 298 to truncate the 666-amino acid long protein at amino acid 315). **(B)** Representative western blot of Spartin protein after gene silencing in hNSCs. First lane: hNSC lysed at day 0; second lane: untransfected hNSCs lysed at day 4; third lane: hNSCs transfected with scramble siRNAs lysed at day 4; forth lane: hNSCs transfected with siRNAs specific to *SPART* transcripts lysed at day4; fifth lane: untransfected hNSCs lysed at day 8; sixth lane: hNSC transfected with scramble siRNAs lysed at day 8; seventh lane: hNSCs transfected with siRNAs specific to *SPART* transcripts lysed at day 8. The image showed the complete absence of Spartin protein after 4 and 8 days of silencing. Gamma tubulin was used as reference. **(C-D)** Quantitative analysis of neurite length and neurite number respectively. *SPART*^892dupA^ mutant cells (n=20) showed a strong increase in the number of neurites per cell and in their length compared to *SPART*^wt^ (n=20). *** p < 0.0001; *p < 0.05, means ± SEM.

**Supplementary Figure 2.**
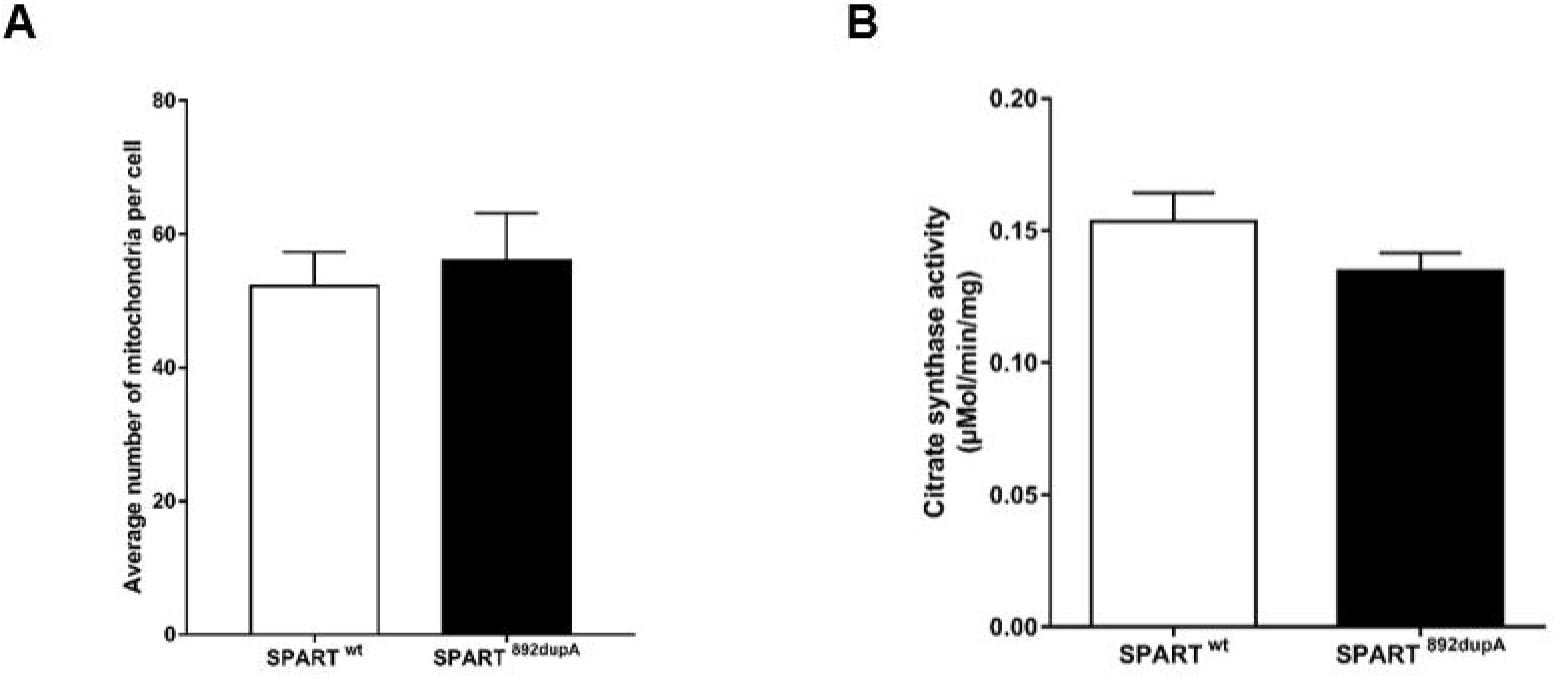
Mitochondrial quantification of mass and citrate synthase activity. Mitochondrial mass measurement with mito-GFP probe in live-cells. Fluorescence signal was quantified using ImageJ standard tools. Error bars indicate ± SD. *, P < 0.01. **(B)** Citrate synthase (CS) activity measurement in *SPART*^wt^ (n=35) and *SPART*^892dupA^ (n=35) cells. Data were normalized on protein content measured by the Lowry assay.

**Supplementary Figure 3:**
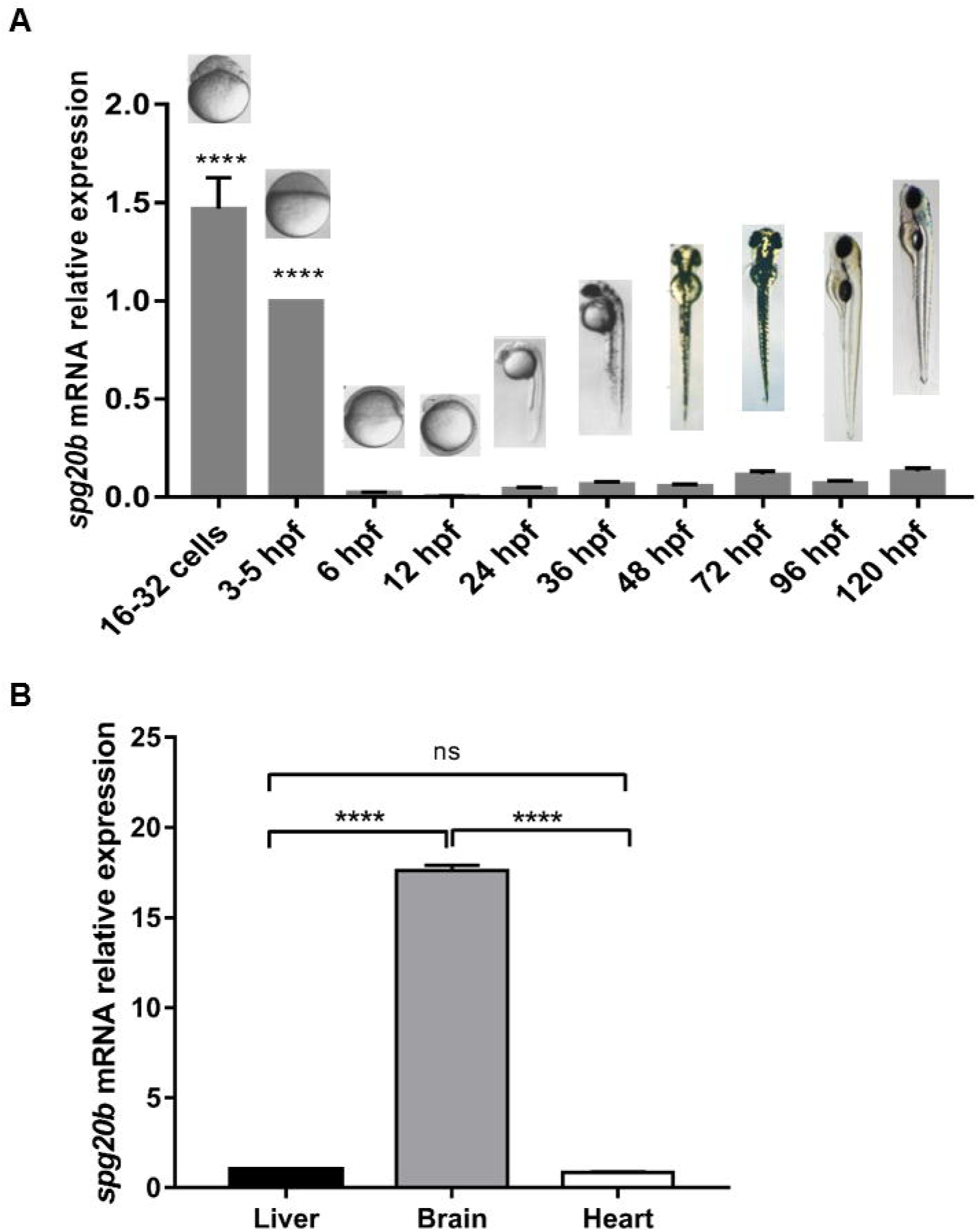
Gene expression analysis in zebrafish. **(A)** Gene expression analysis of *spg20b* gene at different zebrafish developmental stages, spanning from 16–32 cells, up to 120 hours post-fertilization (hpf). A very high *spg20b* expression was identified at the 16–32 cells and 3–5 hpf stages (including cleavage and blastula periods), suggesting an important role for Spartin in these initial developmental phases. Representative optical images of zebrafish embryos/larvae morphology at each development stages are shown. **(D)**. Gene expression analysis of *spg20b* gene in zebrafish adult heart, liver and brain tissues. We observed a *spg20b* expression 18 folds times higher in brain compared to liver and heart. ****P < 0.0001, means ± SEM.

**Supplementary Table 1.**
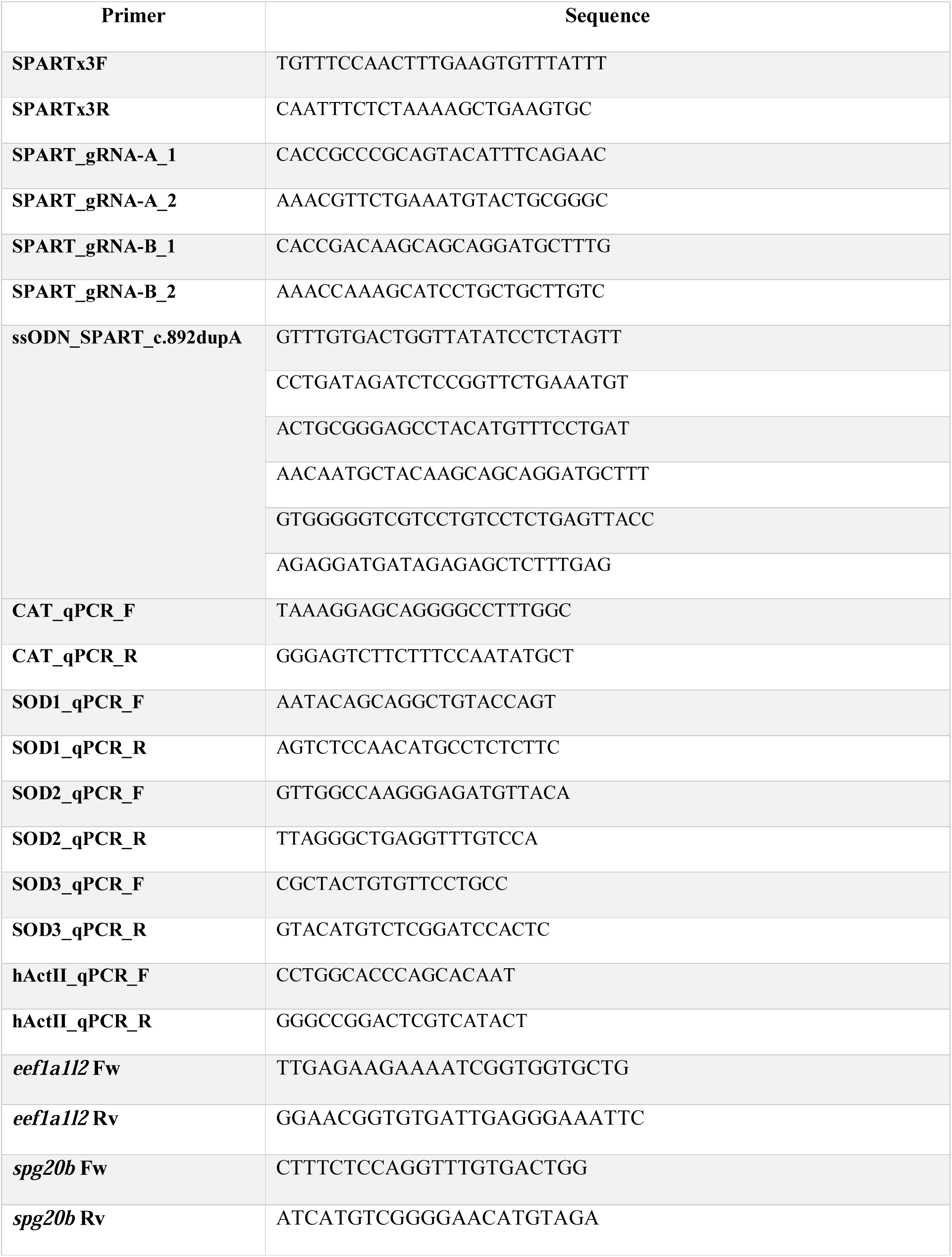
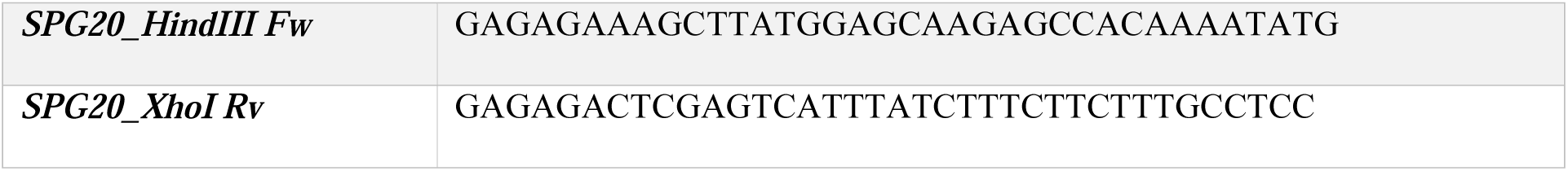
Primer sequences for Sanger method confirmation of WES mutation in *SPART*, gRNA sequences, DNA donor oligo and primers for RT-qPCR are reported.

**Supplementary Table 2.**
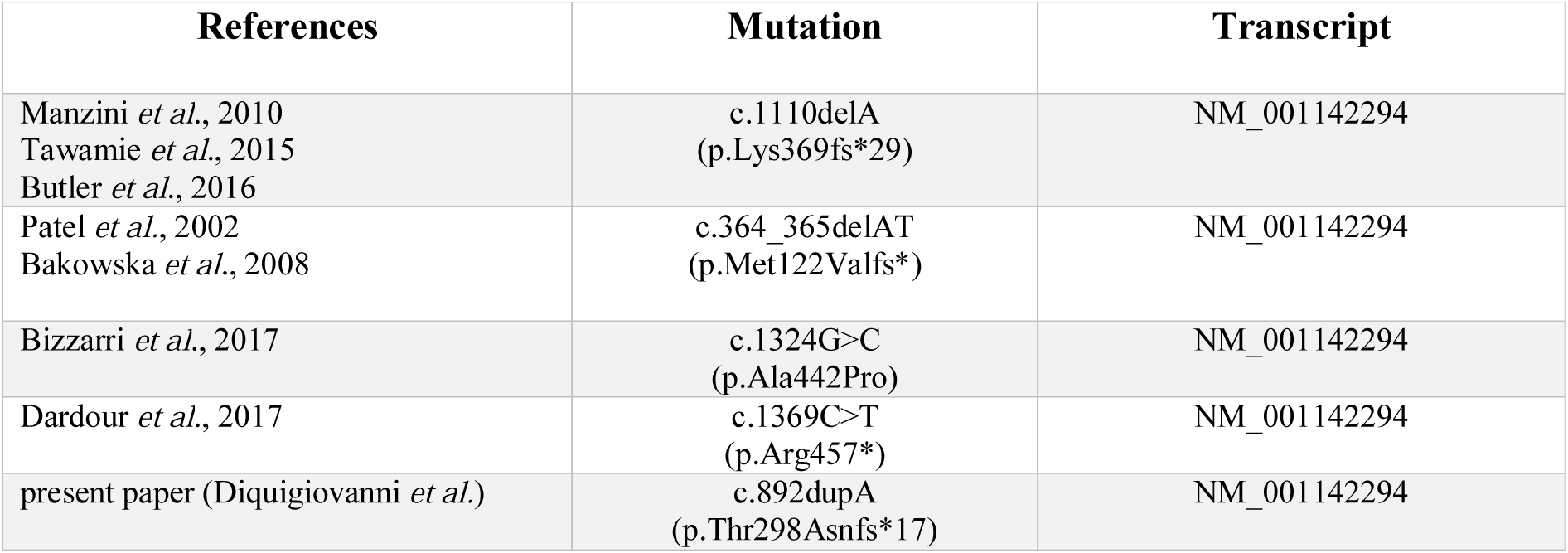
Mutations reported in *SPART* in Troyer syndrome.

