## Supplementary material for "In Troyer syndrome Spartin loss induces Complex I impairments and alters pyruvate metabolism"

**Supplementary Table 1.** Primer sequences for Sanger method confirmation of WES mutation in *SPART*, gRNA sequences, DNA donor oligo and primers for RT-qPCR are reported.

| Primer | Sequence |
| --- | --- |
| SPARTx3F | TGTTTCCAACTTTGAAGTGTTTATTT |
| SPARTx3R | CAATTTCTCTAAAAGCTGAAGTGC |
| SPART_gRNA-A_1 | CACCGCCCGCAGTACATTTCAGAAC |
| SPART_gRNA-A_2 | AAACGTTCTGAAATGTACTGCGGGC |
| SPART_gRNA-B_1 | CACCGACAAGCAGCAGGATGCTTTG |
| SPART_gRNA-B_2 | AAACCAAAGCATCCTGCTGCTTGTC |
| ssODN_SPART_c.892dupA | GTTTGTGACTGGTTATATCCTCTAGTT |
|  | CCTGATAGATCTCCGGTTCTGAAATGT |
|  | ACTGCGGGAGCCTACATGTTTCCTGAT |
|  | AACAATGCTACAAGCAGCAGGATGCTTT |
|  | GTGGGGGTCGTCCTGTCCTCTGAGTTACC |
|  | AGAGGATGATAGAGAGCTCTTTGAG |
| CAT_qPCR_F | TAAAGGAGCAGGGGCCTTTGGC |
| CAT_qPCR_R | GGGAGTCTTCTTTCCAATATGCT |
| SOD1_qPCR_F | AATACAGCAGGCTGTACCAGT |
| SOD1_qPCR_R | AGTCTCCAACATGCCTCTCTTC |
| SOD2_qPCR_F | GTTGGCCAAGGGAGATGTTACA |
| SOD2_qPCR_R | TTAGGGCTGAGGTTTGTCCA |
| SOD3_qPCR_F | CGCTACTGTGTTCCTGCC |
| SOD3_qPCR_R | GTACATGTCTCGGATCCACTC |
| hActII_qPCR_F | CCTGGCACCCAGCACAAT |
| hActII_qPCR_R | GGGCCGGACTCGTCATACT |
| *eef1a1l2* Fw | TTGAGAAGAAAATCGGTGGTGCTG |
| *eef1a1l2* Rv | GGAACGGTGTGATTGAGGGAAATTC |
| *spg20b* Fw | CTTTCTCCAGGTTTGTGACTGG |
| *spg20b* Rv | ATCATGTCGGGGAACATGTAGA |
| *SPG20_HindIII Fw* | GAGAGAAAGCTTATGGAGCAAGAGCCACAAAATATG |
| *SPG20_XhoI Rv* | GAGAGACTCGAGTCATTTATCTTTCTTCTTTGCCTCC |

**Supplementary Table 2.** Mutations reported in *SPART* in Troyer syndrome.

| References | Mutation | Transcript |
| --- | --- | --- |
| Manzini *et al*., 2010  Tawamie *et al*., 2015  Butler *et al*., 2016 | c.1110delA  (p.Lys369fs*29) | NM_001142294 |
| Patel *et al.*, 2002  Bakowska *et al*., 2008 | c.364_365delAT  (p.Met122Valfs*) | NM_001142294 |
| Bizzarri *et al*., 2017 | c.1324G>C  (p.Ala442Pro) | NM_001142294 |
| Dardour *et al*., 2017 | c.1369C>T  (p.Arg457*) | NM_001142294 |
| present paper (Diquigiovanni *et al.*) | c.892dupA  (p.Thr298Asnfs*17) | NM_001142294 |

**Supplementary Figure Legends**

**Supplementary Figure 1**. **Spartin protein analysis in neural cell models. (A)** Representative image of Spartin protein showing the Microtubule Interacting and Trafficking domain (MIT) (aa 16-94) and the senescence domain (aa 427-611). The mutation identified in the two brothers is indicated by an arrow (duplication of nucleotide 892 at amino acid 298 to truncate the 666-amino acid long protein at amino acid 315). **(B)** Representative western blot of Spartin protein after gene silencing in hNSCs. First lane: hNSC lysed at day 0; second lane: untransfected hNSCs lysed at day 4; third lane: hNSCs transfected with scramble siRNAs lysed at day 4; forth lane: hNSCs transfected with siRNAs specific to *SPART* transcripts lysed at day4; fifth lane: untransfected hNSCs lysed at day 8; sixth lane: hNSC transfected with scramble siRNAs lysed at day 8; seventh lane: hNSCs transfected with siRNAs specific to *SPART* transcripts lysed at day 8. The image showed the complete absence of Spartin protein after 4 and 8 days of silencing. Gamma tubulin was used as reference. **(C-D)** Quantitative analysis of neurite length and neurite number respectively. *SPART*^892dupA^ mutant cells (n=20) showed a strong increase in the number of neurites per cell and in their length compared to *SPART*^wt^ (n=20). *** p < 0.0001; *p < 0.05, means ± SEM.

**Supplementary Figure 2. Mitochondrial quantification of mass and citrate synthase activity. (A)** Mitochondrial mass measurement with mito-GFP probe in live-cells. Fluorescence signal was quantified using ImageJ standard tools. Error bars indicate ± SD. *, P < 0.01. **(B)** Citrate synthase (CS) activity measurement in *SPART*^wt^ (n=35) and *SPART*^892dupA^ (n=35) cells. Data were normalized on protein content measured by the Lowry assay.

**Supplementary Figure 3: Gene expression analysis in zebrafish. (A)** Gene expression analysis of *spg20b* gene at different zebrafish developmental stages, spanning from 16–32 cells, up to 120 hours post-fertilization (hpf). A very high *spg20b* expression was identified at the 16–32 cells and 3–5 hpf stages (including cleavage and blastula periods), suggesting an important role for Spartin in these initial developmental phases. Representative optical images of zebrafish embryos/larvae morphology at each development stages are shown. **(D)**. Gene expression analysis of *spg20b* gene in zebrafish adult heart, liver and brain tissues. We observed a *spg20b* expression 18 folds times higher in brain compared to liver and heart. ****P < 0.0001, means ± SEM.
